# Thermodynamic, Kinetic, and Structural Basis of Polyspecificity in the Malaria Sugar Transporter PfHT1

**DOI:** 10.64898/2026.09.13.751325

**Authors:** Arnav Paul, Lingyun Xu, Diwakar Shukla

## Abstract

The malaria parasite Plasmodium falciparum relies on the polyspecific sugar transporter PfHT1 for survival in diverse nutrient environments, making the molecular basis of its polyspecificity a potential avenue for antimalarial drug discovery. However, the structural, thermodynamic and kinetic basis of the emergence of polyspecificity in membrane transporters are not established in literature. Despite advances in AI for protein engineering, the design of selective/promiscuous membrane transporter proteins remains challenging as it requires connecting substrate recognition to the energetics of the transport cycle. Here, we examine the molecular origins of polyspecificity in PfHT1 transporter using molecular dynamics simulations of substrate transport and experimental validation of the key residues that tune specificity. From a structural viewpoint, we hypothesize that polyspecific membrane transporter contains flexible structural elements that could reshape the binding pocket in a substrate-dependent manner. For PfHT1, the cracking of the extracellular helix TM7b allows flexible re-shaping of the binding pocket similar to the promiscuous enzymes. The thermodynamic signature of polyspecificity emerges in the form of the substrate dependent stabilization of transport-competent states. From a kinetic perspective, this stabilization asymmetrically alters transition kinetics and channels the flux through diverse conformational pathways that facilitate the conformational transitions from outward to inward facing states. To investigate the contribution of the underlying interaction network to sugar specificity, we used graph-attention learning to prioritize mutant transporters with distinct substrate preferences. Distinct glucose- and fructose-dependent growth phenotypes identify positions with differential effects on sugar utilization, providing functional evidence relevant to substrate specificity within the polyspecific transporter. Together, these findings support a mechanism in which structural, thermodynamic and kinetics observables connect polyspecificity to conformational energetics, kinetics, and structural plasticity.

## Introduction

Membrane transporters are indispensable for cellular physiology, regulating the controlled movement of nutrients, metabolites, and signaling molecules across biological membranes. Among them, the major facilitator superfamily (MFS) represents one of the largest and most diverse classes of secondary active transporters.^1,2^ Members of this family operate through the canonical rocker-switch alternating access mechanism, in which substrate binding and release are coupled to large-scale conformational transitions.^3,4^ Sugar porter family is a major constituent in the MFS superfamily.^5^ Many structural biochemical studies have defined key conformational states of sugar porters like GLUTs, bacterial sugar transporter in *E. Coli* XylE, *Plasmodium falciparum* sugar transporter PfHT1, plant transporters like STP6, STP10 and SUC1 from *Arabidopsis thaliana*, expanding the understanding of the stable conformations of the transporters. ^6–22^ Membrane transport, however, is not determined solely by the structures of the outward and inward facing states, but by the energetic and kinetic landscape connecting them. Substrate binding can shift conformational equilibria, stabilize intermediate states, and alter the barriers separating functionally relevant conformations, thereby coupling ligand recognition to alternating access. Ligand-dependent remodeling of conformational ensembles has been observed across diverse transporter families, including MFS, LeuT-fold, and MATE transporters, suggesting that modulation of the underlying energy landscape is a general feature of membrane transport.^1,23,24^ However, the kinetic pathways, transition rates and energetic coupling between substrate binding and conformational change remain poorly resolved for sugar porters, particularly from molecular mechanism perspective. Recent studies have explored the molecular mechanism of GLUT5 transport cycle using biased sampling methods and reconstructing the transport cycle for sugar transporters by leveraging evolutionary information.^25,26^

In this context, the Plasmodium falciparum hexose transporter 1 (PfHT1) falls under the promiscuous end of sugar transporters.^27,28^ Here, using PfHT1 as a model membrane sugar transporter, we combine extensive adaptive molecular dynamics simulations with Markov state models, transition-path theory, graph-based machine learning, and functional experiments to determine how substrate binding reorganizes the transport cycle. As the hexose transporter in the malaria parasite, PfHT1 is indispensable for parasite survival throughout its blood-stage lifecycle.^29^ Unlike mammalian GLUT transporters, PfHT1 is notably promiscuous, capable of facilitating the uptake of multiple hexoses and structurally related compounds.^15^ This polyspecificity expands the parasite’s access to alternative carbon sources: fructose supplementation can sustain intraerythrocytic parasite growth under glucose-limited culture conditions, suggesting a potential nutritional advantage when glucose availability declines and alternative sugars are sufficiently abundant.^28^ Promiscuity in substrate recognition is a recurring feature of many MFS transporters. Several mechanisms may enable this functional flexibility: (i) a broad conformational ensemble that accommodates substrates without imposing large energetic penalties, (ii) a distributed and dynamically reorganizing residue interaction network that can maintain substrate engagement across different stages of transport, and (iii) adaptable transport kinetics and conformational pathways that permit productive translocation despite differences in substrate interactions. How these energetic, interaction, and kinetic mechanisms collectively contribute to substrate recognition and transport remains poorly understood.

One structural mechanism that may connect these energetic, interaction, and kinetic layers is helix cracking, in which a localized region of a helix undergoes substantial deformation or partial disruption of its regular helical geometry during a functional conformational transition.^30,31^ Rather than requiring an entire transmembrane helix to move as a rigid body, cracking can localize structural deformation to a short segment, providing a route for larger conformational rearrangements while limiting the accumulation of strain. In sugar porters, TM7b is particularly well positioned to serve this role because it contributes both to the extracellular gate and to the substrate-binding environment. Structural studies of PfHT1 and related GLUT transporters have captured TM7b in sharply bent or broken configurations during occlusion, linking local TM7b deformation to sugar coordination and extracellular-gate closure.^6,15^ Because changes in TM7b geometry simultaneously alter the binding cavity and the accessibility of conformational states, helix cracking provides a potential mechanism connecting binding-pocket plasticity, conformational energetics, and substrate promiscuity.

Understanding PfHT1 is not only a fundamental biophysical question but also a therapeutic opportunity. ^32^ Malaria remains a global health burden, and despite significant progress, targeting nutrient uptake pathways remains underexplored relative to enzyme-based targets, limiting opportunities for selective therapeutic intervention. Nutrient uptake pathways such as PfHT1 represent an attractive but underexploited class of targets.^33^ Structural studies have already highlighted exploitable differences between PfHT1 and human GLUT homologues, and small-molecule inhibitors have been identified that selectively block parasite glucose uptake.^15,16^ However, rational inhibitor design requires a detailed understanding of the conformational cycle, gating elements, and residue-level determinants that underpin transport function, which remains a challenge.

Here, we address this gap by resolving the full conformational cycle of PfHT1 using extensive unbiased all-atom molecular dynamics simulations analyzed through Markov state models (MSMs).^34^ Markov state models provide a principled framework to extract long-timescale kinetics and transition pathways from short timescale simulations, enabling quantitative characterization of transport cycles that are otherwise inaccessible.^34–37^ Transport mechanisms of multiple classes of transporters have been identified using MSMs. ^38–43^ We built atomistic model of PfHT1 in complex *Plasmodium* membrane and ran over 800 *µ*s of MD simulations using adaptive sampling methods. Our results quantify the kinetics and energetics of conformational transitions in apo and glucose-bound PfHT1, revealing how substrate binding reshapes conformational exchange and biases progression through the transport cycle. We identify progressive TM7b helix cracking, quantified through changes in TM7b kink geometry, as a local structural transition coupled to extracellular-gate closure and substrate progression. In addition to state populations and transition times, we resolve the OF-to-IF reactive-path ensemble using transition-path theory and latent-space pathway clustering.^44,45^ This analysis distinguishes conformational heterogeneity in apo PfHT1 from the substrate-induced convergence of glucose-bound pathways through the stabilized OC state. We further examine whether this convergence extends to the substrate-binding pocket. MSM-weighted ensemble analysis shows that apo PfHT1 samples several pocket configurations, whereas glucose stabilizes a restricted pocket geometry coupled to extracellular-gate closure and TM7b cracking.

Recent advances in graph-based machine learning provide an opportunity to extract interpretable residue-level determinants directly from simulation data, yet their application to membrane transport mechanisms remains limited. ^46,47^ To pinpoint residue-level determinants, we develop a graph attention network (GAT) model, which systematically highlighted critical residues for gating and substrate coordination.^48^ These include extracellular gate residues, binding-pocket residues, and intracellular gate residues. We experimentally tested GAT prioritized residues through alanine substitutions and yeast growth assays in glucose and fructose containing media. The widespread growth defects among these variants provide functional support for the transport-associated residue network identified by the model. Together, these results provide a residue-level framework for understanding its promiscuous substrate profile, and increases our understanding of PfHT1 for rational drug discovery against malaria by targeting nutrient uptake.

## Results

### PfHT1 transport cycle in apo and glucose-bound states

The MFS family transporters usually follow a 12 TM helix topology with TM1-TM7 and TM4-TM10 forming the gates. Previous studies on PfHT1 have shown that TM1-TM7 form the extracellular gates and TM4-TM10 form the intracellular gates, in agreement with MFS family transporters. The extracellular and intracellular gating distances serve as a feature to describe the conformations of PfHT1. The outward facing (OF) state is characterized by large extracellular gating distance and small intracellular gating distance and the other way round for the inward facing (IF) state. Occluded (OC) state can be described when both intracellular and extracellular distances are small. PfHT1 gating dynamics can be described by the distance between the N48-S315 distance for the extracellular gates and S153-E417 residue pair distance for the intracellular gates (Figure 1A). All MSM-weighted simulation data has been projected on the gating distance metrics to obtain the conformational landscape plots for apo and holo PfHT1 (Figure 1B,C). Bootstrap-derived uncertainties for the apo and glucose-bound gating free-energy landscapes are shown in Figure S3.

**Figure 1:**
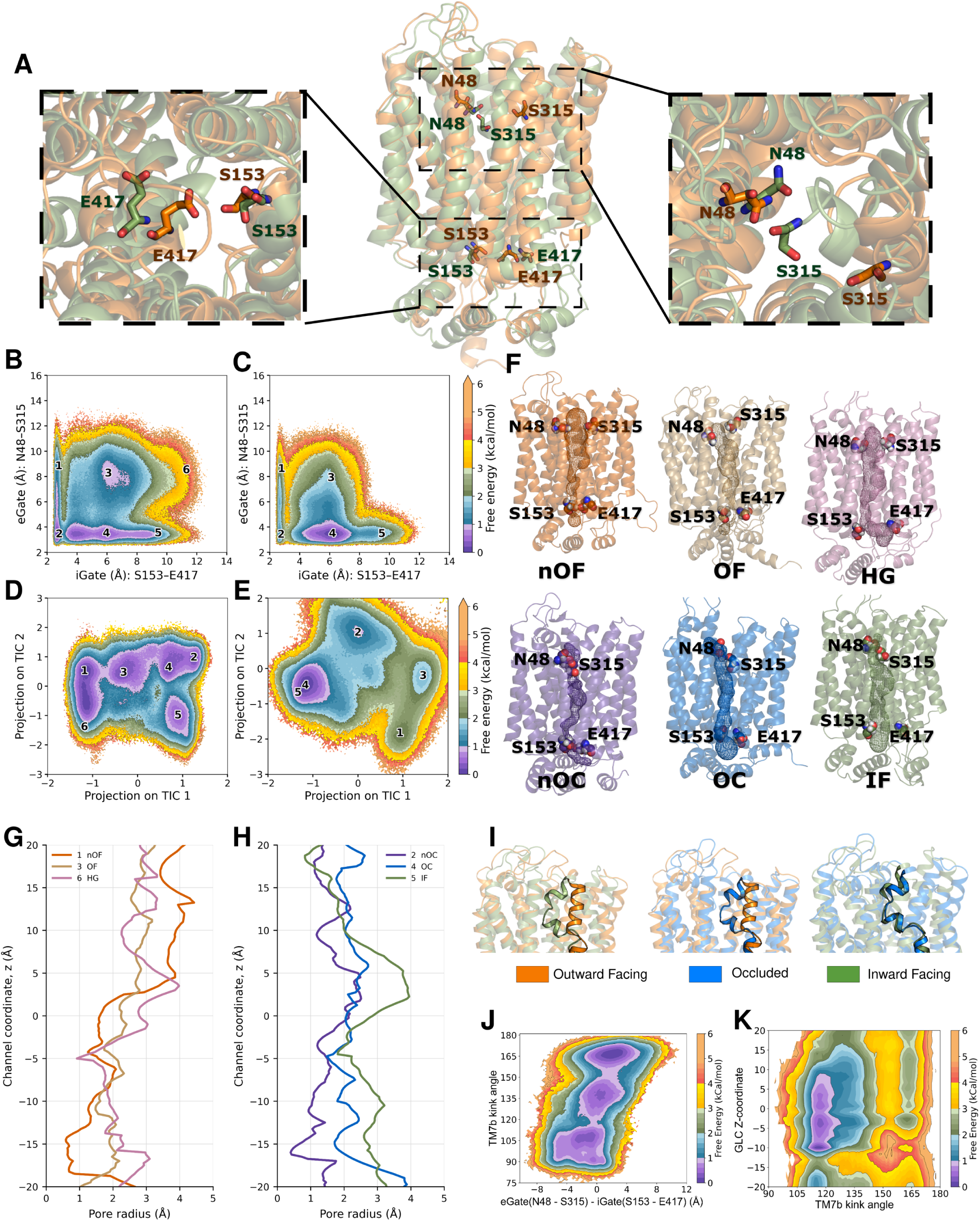
Conformational landscapes, pore geometries, and TM7b coupling in apo and glucose-bound PfHT1. (A) Overlay of representative outward-facing (orange) and inward-facing (green) structures. Enlarged views show the intracellular S153–E417 and extracellular N48–S315 gating pairs. (B,C) MSM-reweighted conformational free-energy landscapes projected onto the extracellular- and intracellular-gate distances for (B) apo and (C) glucose-bound PfHT1. Numbered labels denote narrow outward-facing (nOF, 1), narrow occluded (nOC, 2), outward-facing (OF, 3), occluded (OC, 4), inward-facing (IF, 5), and hourglass (HG, 6) states; a distinct HG state is observed only in the apo ensemble. (D,E) Corresponding free-energy landscapes projected onto the first two time-lagged independent components for apo (D) and glucose-bound (E) PfHT1, with macrostate positions indicated. (F) Representative structures of the six apo macrostates showing the transport tunnel and gating residues. (G,H) Pore-radius profiles along the channel coordinate for (G) nOF, OF, and HG and for (H) nOC, OC, and IF. (I) Pairwise structural comparisons of TM7b in outward-facing (OF, orange), occluded (OC, blue), and inward-facing (IF, green) conformations, highlighting changes in TM7b geometry across the transport cycle. (J) MSM-reweighted apo free-energy landscape projected onto the TM7b kink angle and the difference between the extracellular and intracellular-gate distances, *d*_N48*−*S315_ *− d*_S153*−*E417_. (K) Glucose-bound free-energy landscape projected onto the glucose z-coordinate and TM7b kink angle. Positive z-values point toward the extracellular side and negative values toward the intracellular side.

Six apo macrostates were identified: narrow outward-facing (nOF; state 1), narrow occluded (nOC; state 2), outward-facing (OF; state 3), occluded (OC; state 4), inward-facing (IF; state 5), and hourglass (HG; state 6). In apo PfHT1, the principal conformational route connects OF to OC and then IF, while nOF connects to the more tightly occluded nOC state before reaching IF (Figure 1B). The apparent barriers separating the dominant basins are approximately 1–1.2 kcal/mol in the projected landscapes. Because these values are obtained from low-dimensional projections, they represent projected free-energy barriers rather than direct kinetic activation barriers. The glucose-bound landscape retains the OF–OC–IF sequence but changes the relative stability of the conformational basins (Figure 1C). The nOF and nOC states lie at higher free energy than the connected low-energy region containing OF, OC, and IF, consistent with glucose favoring conformations along the productive transport pathway. A distinct HG macrostate was not resolved in the glucose-bound ensemble. Projection onto the first two time-lagged independent components provides a complementary representation of the slow conformational coordinates used to construct the MSMs (Figures 1D,E). Time-lagged independent component analysis (tICA) identifies combinations of structural features that capture the slowest conformational motions sampled by the transporter. Consistent with the gating-coordinate landscapes, the tICA projections resolve distinct conformational basins separated by modest projected free-energy barriers. In the apo ensemble, transitions between the major basins involve barriers of approximately 1–1.2 kcal/mol, whereas the glucose-bound landscape exhibits a more continuous low-energy pathway with barriers of roughly 1 kcal/mol between the principal states, consistent with substrate binding facilitating conformational exchange. All six apo macrostates can be located in the apo tIC landscape, whereas the glucose-bound tIC landscape contains states 1–5 without a corresponding HG basin. The agreement between the gating coordinate landscape and tIC representations indicates that the principal state assignments are retained across two different low-dimensional descriptions. Projection of both ensembles onto the apo-derived tICA basis provides an additional common-coordinate comparison (Figure S5), with bootstrap-derived uncertainties shown in Figure S6. Mapping the gate distances onto tICA space further shows that tIC1 primarily tracks extracellular gating, whereas tIC2 captures intracellular gating in both systems (Figures S7 and S8).

Representative structures were selected by clustering the simulation ensemble in the first two tICA dimensions (tIC1 and tIC2), with structures corresponding to the identified conformational clusters used to represent each macrostate (Figure 1F). The structures show the N48–S315 and S153–E417 gating pairs together with the accessible tunnel for each of the six apo macrostates. Pore-radius profiles quantify these changes for nOF, OF, and HG (Figure 1G) and for nOC, OC, and IF (Figure 1H). Pore-radius calculation for the OF, OC and IF ensembles from the simulations are shown in Figure S1. The HG state combines access from both sides with a constriction near the center of the transporter, distinguishing it from the productive OC intermediate despite having relatively open extracellular and intracellular gates. The absence of a distinct HG basin in both representations of glucose-bound PfHT1 is consistent with substrate binding not favoring the centrally constricted geometry (Figures 1C,E). In the absence of substrate-mediated interactions, apo PfHT1 samples a broader conformational ensemble, including conformations that are not appreciably populated in the glucose-bound landscape. In contrast, glucose-mediated interactions preferentially stabilize specific conformations along the transport cycle, resulting in a more restricted conformational ensemble. This redistribution provides a thermodynamic starting point for understanding polyspecificity, PfHT1 accesses multiple conformational states, and substrate interactions change their relative stability. We next examined the local structural changes that could couple this stabilization to productive gating.

### TM7b helix cracking couples substrate progression to gate closure

Large-scale alternating access requires substantial rearrangement of transmembrane helices, but these motions need not occur through rigid-body displacement of entire helices. Localized deformation of secondary structure, often described as helix cracking, can concentrate conformational change within a short segment of a helix. In PfHT1, TM7b undergoes such a transition across the transport cycle. The helix is straight in the OF state but adopts progressively more sharply bent, or cracked, geometries in the OC and IF states, with the local deformation centered near S315 and N316 (Figure 1I). We therefore use the TM7b kink angle as a geometric measure of the extent of helix cracking. To quantify this coupling, the MSM-reweighted apo ensemble was projected onto the TM7b kink angle and the difference between the extracellular and intracellular-gate distances, Δ*d*_gate_ = *d*_N48*−*S315_ *− d*_S153*−*E417_ (Figure 1J). The landscape contains regions corresponding to the relatively straight OF geometry (*∼*170*^◦^*), the intermediately bent OC geometry (*∼*135*^◦^*), and the more strongly bent IF ensemble (*∼*90–120*^◦^*), demonstrating progressive TM7b bending along the OF-to-IF conformational transition. Thus, TM7b bending is correlated with progression from outward to inward access. We also projected the TM7b kink angle on the gating landscapes and observed a clear correlation with the extracellular gating distances in both apo and holo systems (Figure S2). Sugar transporter family GLUTs and other MFS transporters also share a similar feature, however the mechanistic reason for this kink in TM7b was unknown. A recent study by Drew and coworkers proposed a two-step mechanism for sugar translocation in which TM7b kinking is a key structural transition that facilitates substrate movement through the transporter. This experimentally derived mechanism provides independent support for our simulations, which identify progressive TM7b bending as a major conformational change coupled to gate closure and glucose translocation in PfHT1.^17^

We next projected the glucose z-coordinate against the TM7b kink angle to determine how substrate position is associated with helix geometry (Figure 1K). The z-coordinate of the glucose provides an interpretable metric to understand the progress of substrate transport. Positive z-values correspond to the extracellular side, values near the center describe the binding-pocket region, and negative values correspond to the intracellular side. The low free energy regions vary with glucose position, demonstrating coupling between substrate progression and TM7b geometry. Extracellular and binding-pocket glucose positions are associated with an OF to OC like TM7b geometry, whereas intracellular positions are associated with more bent, IF-like configurations. Notably, the highest free-energy region occurs near a TM7b angle of 165° when glucose has progressed toward the intracellular side (z = −10 Å), indicating that deep substrate translocation is incompatible with a fully open, OF-like TM7b geometry. This suggests a conformational-selection mechanism in which glucose can initially enter the transporter when TM7b is straight and the extracellular gate is open, but progression toward the OC and IF states requires substrate engagement with conformations that permit TM7b bending and subsequent gate closure. Thus, substrate entry can occur while TM7b remains comparatively uncracked, whereas progression toward the OC and IF ensembles is associated with increasing local helix cracking and extracellular-gate closure. These results support a model in which TM7b bending helps close the extracellular side after substrate entry and accompanies opening toward the cytoplasm. Bootstrap-derived uncertainties for the landscapes relating TM7b bending to conformational change and glucose position are shown in Figure S4. Because TM7b contributes to both the substrate environment and extracellular gating, its flexibility offers a means of accommodating changes in substrate coordination while preserving alternating access. This coupling provides a candidate structural basis for polyspecificity, but its functional consequences also depend on how substrate binding changes the rates of conformational exchange.

### Kinetics of conformational change and glucose transport for PfHT1

To resolve the kinetics of the transport cycle, we calculated mean first passage times (MF-PTs) between the major conformational states of PfHT1 in both the apo and holo cycles. These calculations provide quantitative estimates of the relative accessibility of outward-facing (OF), occluded (OC), inward-facing (IF), and transient states during the transport process. To determine how substrate binding reshapes PfHT1 kinetics, we compared MF-PTs for directed transitions between OF, OC, and IF in the apo and glucose-bound models (Figure 2A). In the comparison presented in panel A, glucose accelerated OF to OC, OC to IF, and overall OF to IF by 2.88, 4.45, and 4.25 fold, respectively. The reverse transitions changed asymmetrically: IF to OC was 2.96-fold faster in the glucose-bound model, whereas OC to OF and overall IF to OF were 1.43 and 1.42-fold slower. Glucose therefore does not uniformly accelerate every transition; it preferentially promotes progression toward IF while suppressing return toward OF. Directional bias was quantified using the rectification index

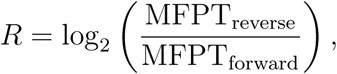

where positive values indicate a preference for the left-to-right direction in the state-pair label (Figure 2B). Apo PfHT1 showed mixed directionality, with *R* = *−*0.24 for OF-OC, *R* = *−*0.48 for OC-IF, and *R* = *−*0.71 for the overall OF-IF transition. In contrast, glucose-bound PfHT1 showed strong forward rectification for OF-OC (*R* = +1.81) and overall OF-IF (*R* = +1.89), while OC-IF was nearly reciprocal (*R* = +0.11). Substrate binding therefore reverses the overall OF-to-IF kinetic bias, producing a strong asymmetry favoring inward-facing progression under the modeled conditions. Here, rectification refers to asymmetry in conditional MFPTs and does not imply a nonequilibrium net flux.

**Figure 2:**
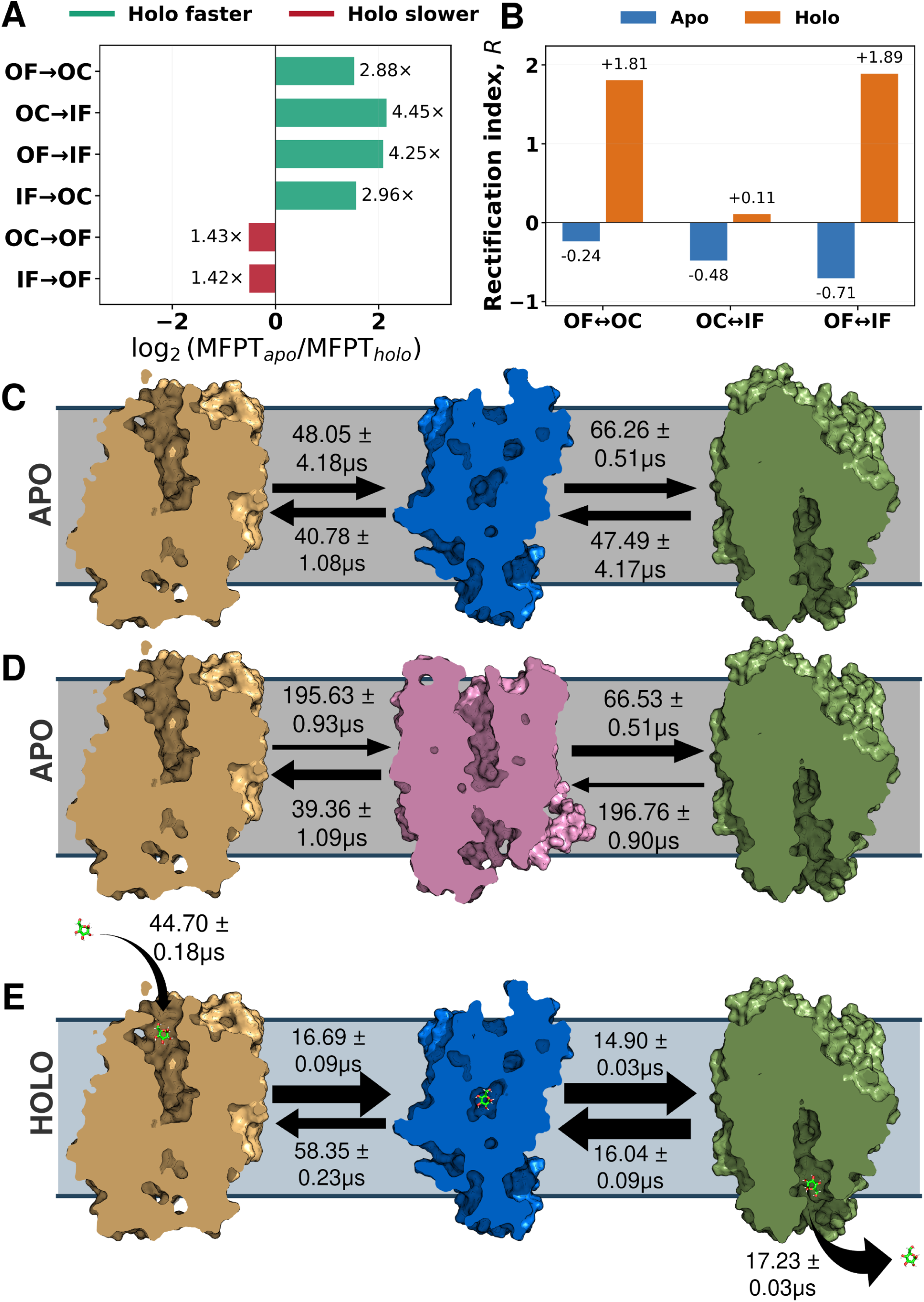
Glucose binding asymmetrically remodels PfHT1 conformational kinetics. (A) Comparison of apo and glucose-bound MFPTs for the indicated directed transitions. Bar positions show log_2_(MFPT*_apo_/*MFPT*_holo_*). (B) Rectification index *R* = log_2_(MFPT*_reverse_/*MFPT*_forward_*) for the OF-OC, OC-IF, and overall OF-IF transitions in apo and glucose-bound PfHT1. Forward is defined as the left-to-right direction in each label; *R >* 0 indicates forward bias and *R <* 0 indicates reverse bias. (C,D) Apo MFPTs for transitions through the primary OC intermediate (C) and the kinetically disfavored HG intermediate (D). (E) Glucose-bound cycle showing glucose entry into the outward-facing cavity, conformational exchange among OF, OC, and IF, and glucose release from the inward-facing state. Tan, blue, pink, and green structures denote OF, OC, HG, and IF, respectively.

Route-resolved MFPTs distinguish the principal OC-mediated apo pathway from the kinetically disfavored HG pathway (Figures 2C,D). Along the OC-mediated route, the OF to OC and OC to IF steps had MFPTs of 48.05 *±* 4.18 and 66.26 *±* 0.51 *µ*s, respectively. The corresponding reverse IF to OC and OC to OF steps had MFPTs of 47.49 *±* 4.17 and 40.78 *±* 1.08 *µ*s respectively (Figure 2C).

Entry into HG has substantially slower MFPTs, OF to HG and IF to HG required 195.63 *±* 0.93 and 196.76 *±* 0.90 *µ*s, respectively. Escape from HG was faster, with HG to OF and HG to IF MFPTs of 39.36 *±* 1.09 and 66.53 *±* 0.51 *µ*s (Figure 2D). Entry into HG, rather than every transition involving HG, is therefore kinetically unfavorable, explaining why HG forms an unproductive alternative to the primary OC-mediated route.

In the glucose-bound cycle, glucose entered the outward-facing cavity with an MFPT of 44.70 *±* 0.18 *µ*s (Figure 2E). The subsequent OF to OC and OC to IF steps occurred in 16.69 *±* 0.09 and 14.90 *±* 0.03 *µ*s, respectively. The reverse OC to OF transition was substantially slower at 58.35 *±* 0.23 *µ*s, whereas IF to OC remained similar to the forward OC to IF step at 16.04 *±* 0.09 *µ*s. Glucose release from the inward-facing cavity occurred in 17.23 *±* 0.03 *µ*s. Substrate binding therefore remodels the transition kinetics asymmetrically, it accelerates progression toward IF while slowing key reverse transitions toward OF. This kinetic rectification funnels glucose-bound PfHT1 through the OC-mediated pathway without implying that every transition in the cycle is accelerated. These results extend the recognition problem from substrate accommodation to the kinetics of productive transport. For polyspecificity, the relevant question is therefore whether different sugars can promote the conformational transitions needed for translocation, in addition to occupying the binding pocket.

### Transition-tube analysis reveals heterogeneous apo pathways and glucose-induced channeling through the OC state

To characterize how PfHT1 transitions from the OF to IF state, we applied transition-path theory (TPT) to the apo and glucose-bound MSMs. While the MSM kinetics quantify the timescales of transitions between conformational states, TPT resolves the ensemble of reactive pathways that connect defined OF and IF end states through the underlying microstate network. Because the resulting reactive ensemble contains many individual microstate sequences, we further applied VAE-based pathway clustering to group geometrically similar paths into broader transition tubes. The 20,000 highest-flux paths accounted for 89.6% and 94.5% of the total reactive current in the apo and glucose-bound systems, respectively (Figure S19), capturing the majority of the OF-to-IF reactive ensemble in both systems.

K-means clustering partitioned the apo TPT paths in the two-dimensional VAE latent space into two transition tubes carrying 53.5% and 46.5% of the represented path flux (Figure 3A,B). Transition tube 1 initially sampled widening of the extracellular gate from nOF toward OF before proceeding through OC to IF. Transition tube 2 followed a more direct nOF-nOC-IF corridor. These differences were clearer in tICA space than in the two gate distances (Figure S21). The forward committor (*q*^+^) provides a measure of progress along the reactive transition, defined as the probability that a microstate will reach the IF state before returning to the OF state. The committor projections showed that the two tubes traverse different conformational regions while progressing from OF (*q*^+^ = 0) to IF (*q*^+^ = 1) (Figure 3E,G). We next examined the order of the two major gating events along each reactive path, specifically whether extracellular-gate (eGate) closure occurred before intracellular-gate (iGate) opening or vice versa. Neither apo tube corresponded exclusively to a single gate-order mechanism. eGate closure preceded iGate opening for 74.6% and 59.9% of the flux in transition tubes 1 and 2, respectively, whereas iGate opening occurred first for 21.6% and 29.8%. The remaining paths exhibited concerted gate rearrangements. Thus, apo PfHT1 accesses two geometrically distinguishable but mechanistically mixed transition tubes. The contribution from iGate-opening-first configurations remains consistent with the low equilibrium population of HG because a short-lived state can carry reactive current without forming a stable free-energy minimum.

**Figure 3:**
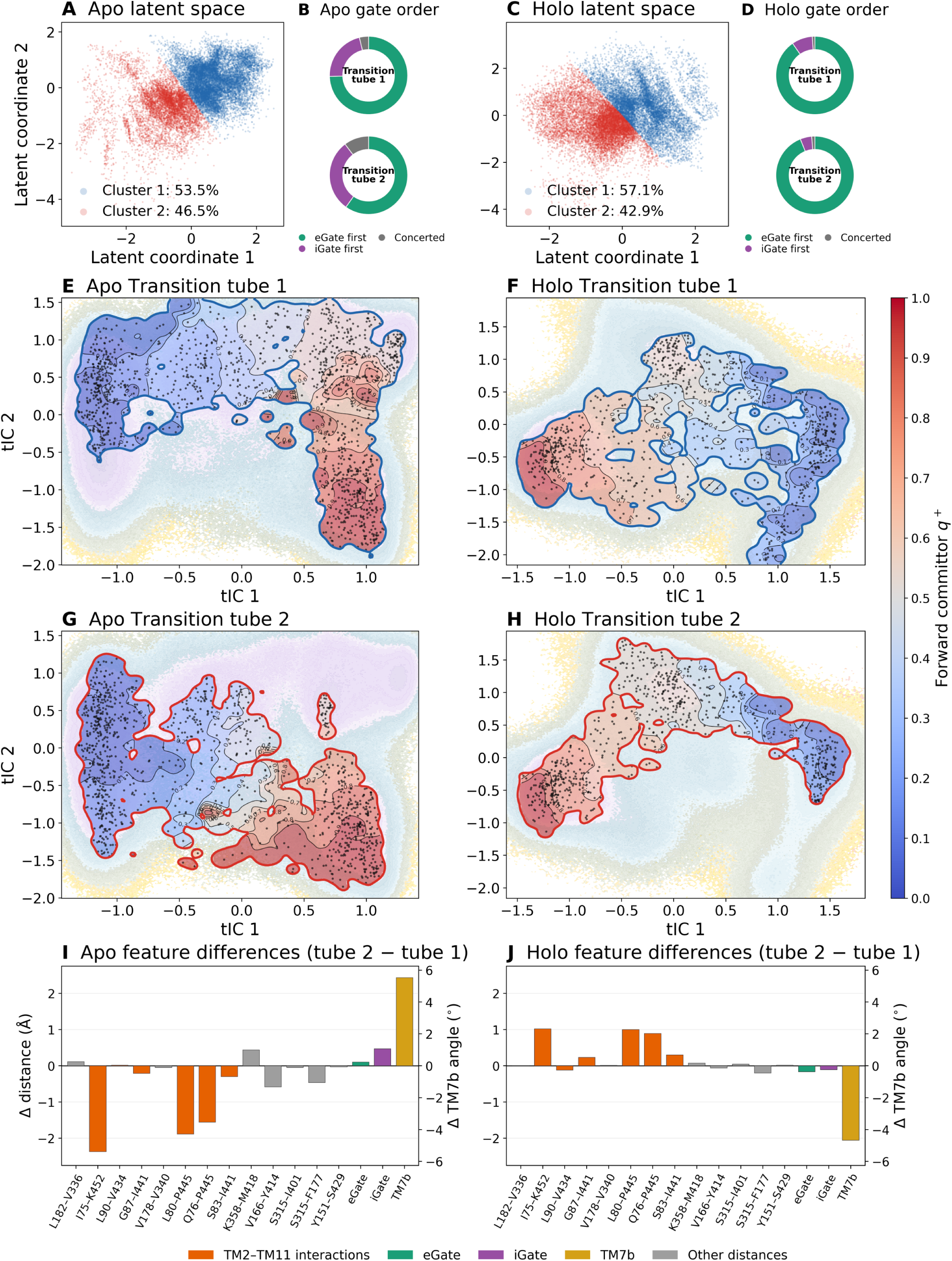
Flux-weighted transition-tube analysis of apo and glucose-bound PfHT1. Two-dimensional VAE embeddings colored by the two K-means clusters for (A) apo and (C) holo. Flux-weighted gate-event order within each cluster for (B) apo and (D) holo PfHT1. Forward-committor projections for transition tube 1 in (E) apo and (F) glucose-bound PfHT1 and transition tube 2 in (G) apo and (H) glucose-bound PfHT1. Black points indicate participating microstates, and committor contours are shown at intervals of 0.1 from OF (*q*^+^ = 0) to IF (*q*^+^ = 1). (I,J) Differences in mean structural features between transition tubes, calculated as tube 2 minus tube 1, for (I) apo and (J) glucose-bound PfHT1.

The glucose-bound TPT paths were similarly projected onto the two-dimensional VAE latent space and separated into two geometric partitions by K-means clustering, carrying 57.1% and 42.9% of the represented path flux, respectively (Figure 3C,D). Both were strongly dominated by eGate closure before iGate opening, with corresponding fractions of 90.3% and 93.9%. Their transition tubes and committor progression converged through the OC region before reaching IF (Figure 3F,H). This convergence is consistent with the pronounced OC minimum in the glucose-bound free-energy landscape. The unrestricted gap statistic favored one diffuse glucose-bound pathway population (Figure S20), the two displayed tubes should therefore be interpreted as geometric subdivisions of a common OC-mediated mechanism rather than independent transport mechanisms.

Tube-level feature differences identify the structural coordinates that distinguish the reactive corridors (Figures 3I,J; Table S5). In apo PfHT1, several TM2–TM11-side distances were approximately 1.6–2.4 Å shorter in tube 2 than in tube 1, whereas the mean extracellular and intracellular-gate distances differed by less than 0.5 Å (Figure 3I). In glucose-bound PfHT1, tube 2 had selected TM2–TM11-side distances approximately 0.9–1.0 Å larger than tube 1 and a TM7b angle approximately 4.7° lower, corresponding to a more strongly kinked configuration under the angle convention used here. Both gate distances differed by less than 0.2 Å (Figure 3J). The transition tubes are therefore distinguished primarily by internal helix packing and local TM7b geometry rather than large differences in global gate distances. The convergence of glucose-bound pathways shows how substrate engagement can organize a heterogeneous conformational ensemble into a common transport mechanism. This suggests that polyspecificity need not require a separate pathway for each sugar, different interaction patterns could instead support progression through shared transport intermediates.

### Contact analysis and graph-attention model identify transport associated residues

To probe the molecular determinants of conformational change and glucose recognition, residue–glucose contacts were quantified using GetContacts package across the MSM-reweighted transport ensemble. The results were organized into three structural regions: the extracellular gating region, the central binding pocket, and the intracellular gating region. Representative snapshots further illustrate residue–glucose interactions at each stage of the cycle (Figure 4A), while the corresponding contact frequencies identify the most frequently engaged residues (Figure 4B). On the extracellular side, contacts are primarily formed by residues from TM7 and TM1, including Asn311, Gln306, Val44, and Ser315. These residues form transient polar and hydrophobic interactions with glucose during the open-to-occluded transition. Within the binding pocket, the most frequent and persistent contacts involve aromatic and polar residues such as Trp412, Asn435, and Trp436, which maintain persistent contacts with glucose throughout the cycle. Additional strong interactions with Thr145, Gln169, and Ser302 provide a complementary polar environment for glucose coordination. The persistence of several of these interactions across a broad range of substrate positions suggests that glucose remains engaged with the central binding region as it progresses toward the intracellular side.

**Figure 4:**
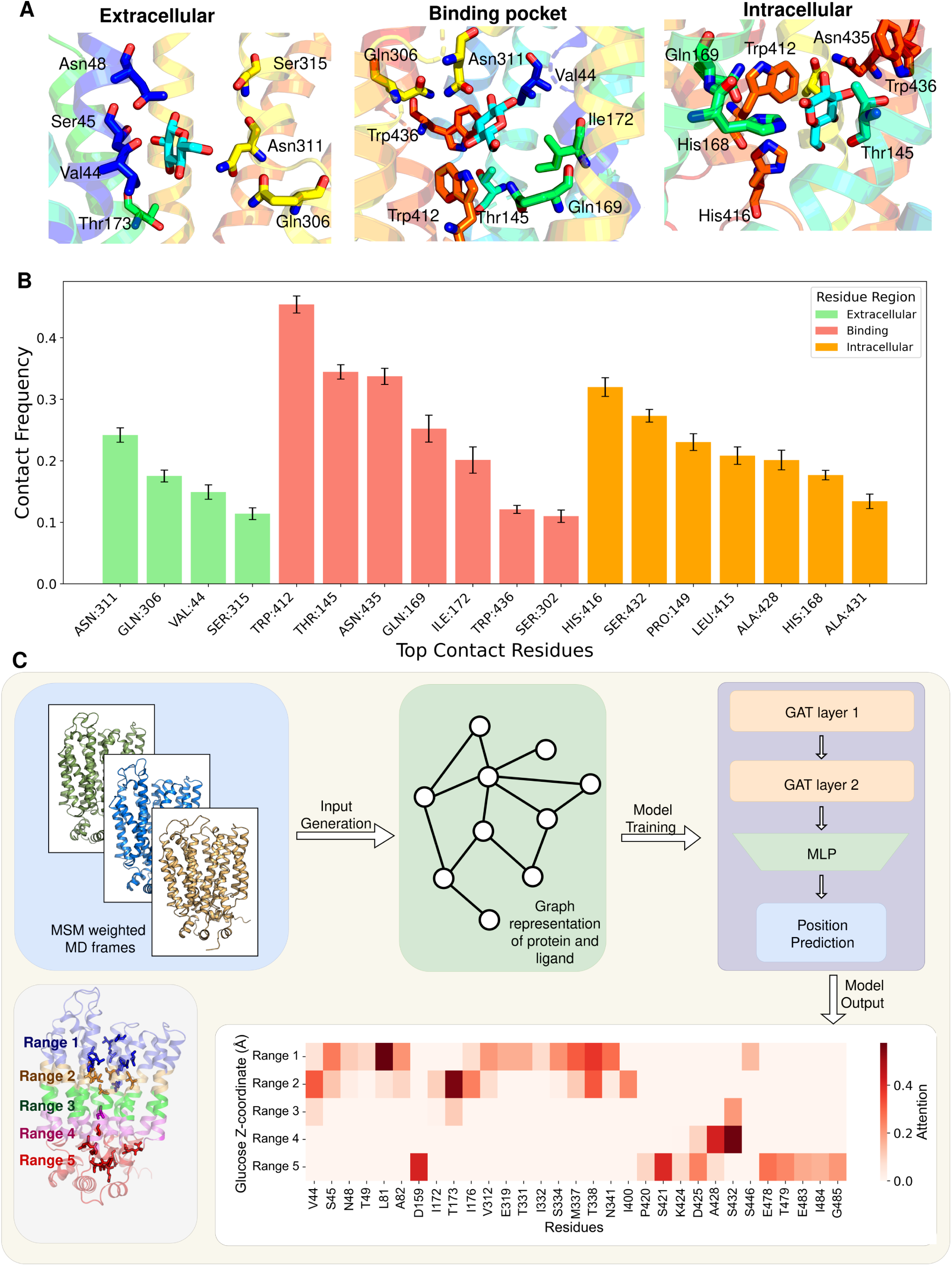
Residue–glucose contact and graph-attention network analysis of PfHT1 transport. (A) Representative residue–glucose contacts in the extracellular region, central binding pocket, and intracellular region. (B) Contact frequencies of the top glucose-interacting residues, colored according to their assigned structural regions. (C) Schematic workflow for identifying residues important for transport using Graph Attention model. Schematic shows the MSM weighted MD frames used to generate graphical input for the GAT model. The output from the GAT is binned based on position of the ligand to identify important residues based on their attention. The PfHT1 structure indicates the corresponding spatial ranges, and the heatmap shows position-dependent attention profiles for selected residues.

As glucose approaches the intracellular vestibule, the interaction profile shifts toward residues including His416, Ser432, Leu415, Pro149, and Ala431. These position-dependent changes accompany substrate movement toward the cytosolic side. Glucose orientation also varies across different regions of the transporter, suggesting that its reorientation is associated with changes in the surrounding interaction environment (Figure S9).

Overall, the contact analysis defines the changing physical interaction environment experienced by glucose as it moves from the extracellular vestibule through the central binding pocket to the intracellular side. These direct residue–glucose contacts provide a structural reference for the Graph attention network based analysis below, which asks a different question: which residues are most informative of substrate position and transport-state progression, including residues whose importance may not arise from persistent direct contact with glucose.

While the contact analysis in Figure 4A,B identifies residues that directly interact with glucose, it does not capture the broader protein environment associated with substrate progression through the transport cycle. We therefore developed a graph attention network (GAT) using MSM-weighted MD frames of the protein–substrate complex to identify residues whose structural environment is informative of glucose position. After training, residue attention scores were grouped according to glucose z-coordinate to determine how the learned attention profile changes as substrate progresses through the transporter. Residues with uniformly high attention across the entire transport process were excluded, as these signals may reflect general structural features rather than position-dependent contributions to transport. We instead focused on residues whose attention changed substantially with glucose position, revealing a dynamic redistribution of residue importance across the transport cycle (Figure 4C).

In the outward-facing ensemble, residues with high attention scores are concentrated primarily in the extracellular region of PfHT1. As glucose approaches the central binding pocket, the distribution of attention shifts away from the extracellular side and toward residues surrounding the substrate-access pathway. These position-dependent changes indicate that different structural features become informative as glucose progresses from the extracellular vestibule toward the center of the transporter.

Within the central binding region, high attention is observed for several polar and aromatic residues located near the glucose-binding pocket. Importantly, this persistence in GAT attention should be interpreted separately from direct glucose-contact persistence identified in Figure 4B. As glucose progresses toward the inward-facing ensemble, attention shifts toward residues on the intracellular side of the transporter. These state-dependent changes indicate that the structural features most informative of glucose position move progressively from the extracellular region, through the central pocket, and toward the intracellular region over the course of the transport cycle.

Several GAT-prioritized positions, including Val44, Ile172, and Ser432, also appear among the prominent glucose-contacting residues identified in Figure 4B, whereas other high attention positions do not. Thus, GAT attention is related to, but not equivalent to, direct contact frequency: the contact analysis reports physical interactions with glucose, whereas the GAT identifies residues whose local structural environment contributes to distinguishing substrate position along the transport pathway. The overlap between the two analyses highlights positions supported by both direct interaction and learned structural information, while non-overlapping positions may reflect more distributed conformational features captured by the model. Several GAT-prioritized residues coincide with functionally or structurally characterized positions in previous PfHT1 studies. Notably, N341 has been shown experimentally to be critical for glucose and fructose transport, while V44, N48, L81, I172, T173, I176, V312, N341, and S446 lie within the substrate/inhibitor interaction region identified from glucose and C3361 bound PfHT1 structures.^16,17,49^ This agreement with independent structural and functional studies supports the biological relevance of the learned attention profile. These analyses identify candidate residues through which substrate engagement could be coupled to the conformational stabilization and pathway convergence described above. If individual positions contribute differently to transport of different sugars, their substitution should produce substrate-dependent functional effects, motivating comparison of glucose and fructose-supported growth.

### Functional testing of computationally prioritized residues by yeast growth assay

To test the functional relevance of the computationally prioritized positions, 26 residues were individually substituted with alanine and the resulting PfHT1 variants were evaluated by yeast growth in glucose and fructose containing media (Figure 5A,B). Growth was quantified using the apparent 0–20 h specific growth rate and normalized independently for each carbon source relative to pooled WT and empty-vector controls.

**Figure 5:**
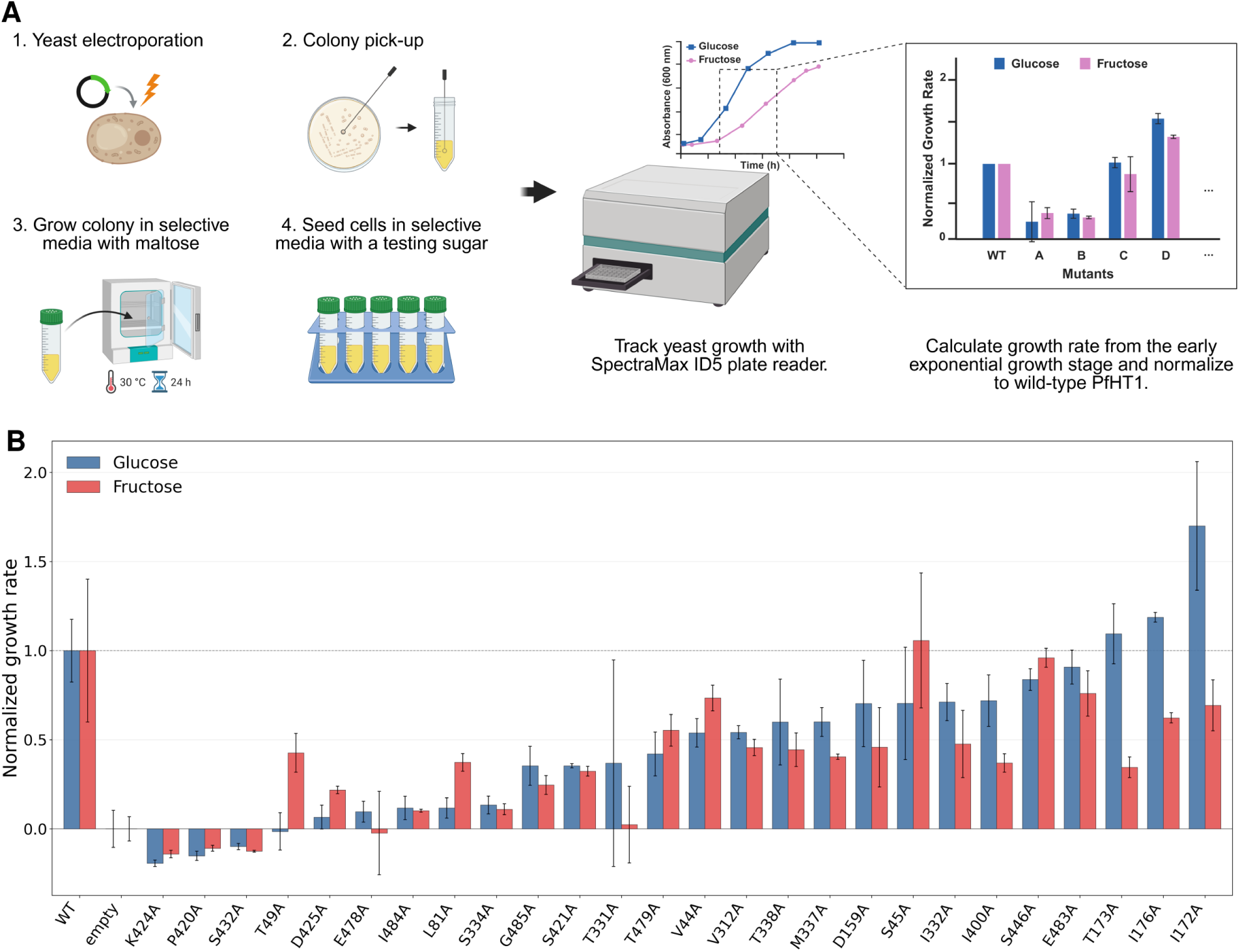
Functional assessment of GAT-selected PfHT1 residues. (A) Schematic experimental workflow for measuring the growth rate of PfHT1 and designed mutants. (B) Apparent 0–20 h growth rates for yeast expressing WT PfHT1, empty vector, or the indicated PfHT1 alanine-substitution variants in glucose- or fructose-containing medium. Bars show mean ± SD. Mutants have *n* = 4 replicate measurements per carbon source; WT and empty-vector controls each have *n* = 12. Glucose, blue; fructose, red.

Alanine substitution produced widespread growth defects, with 23 of 26 variants showing reduced growth relative to WT in glucose, 25 of 26 in fructose, and 22 showing reduced growth under both conditions. P420A, K424A, and S432A produced the strongest phenotypes, with mean growth at or below the empty-vector reference in both media. In contrast, several substitutions showed carbon source dependent effects. T173A, I176A, and I172A retained or exceeded WT-normalized growth in glucose but showed reduced growth in fructose, whereas S45A retained a near-WT fructose phenotype while showing lower growth in glucose. These differences suggest that some prioritized positions may contribute differently to PfHT1-dependent utilization of glucose and fructose.

The prevalence of growth defects among the selected variants provides experimental support for the functional relevance of the residue network identified by the GAT model. The particularly strong effects of several substitutions, together with the substrate-dependent phenotypes of others, further indicate that the prioritized residues do not contribute uniformly to PfHT1 function. However, yeast growth represents an integrated cellular readout that can be influenced by transporter abundance, localization, stability, and substrate uptake. The observed phenotypes therefore support impaired PfHT1-dependent sugar utilization but do not by themselves establish direct changes in substrate binding or transport kinetics. The contrasting glucose and fructose phenotypes connect the glucose-derived computational predictions to the utilization of a second substrate. They support a model in which PfHT1 polyspecificity involves overlapping residue networks whose individual contributions depend on the transported sugar.

### Glucose stabilizes a restricted, transport-competent binding-pocket ensemble

The substrate-dependent growth phenotypes raise the question of how a shared transporter architecture can accommodate different sugars. We therefore compared apo and glucose-bound pocket ensembles to determine whether the conformational restriction observed in the transport landscape and reactive pathways also extends to binding-pocket geometry. Pocket conformations were projected onto principal components derived from the apo ensemble and separated into three apo-defined clusters. In the absence of substrate, PfHT1 broadly populated all three clusters, with occupancies of 23.4%, 41.2%, and 35.5% (Figure 6A,B). The apo transporter therefore accesses a diverse collection of pocket geometries rather than adopting a single dominant binding-site configuration. In contrast, 95.0% of the glucose-bound frames occupied cluster 1. Glucose binding thus markedly restricts the conformational heterogeneity of the binding pocket.

**Figure 6:**
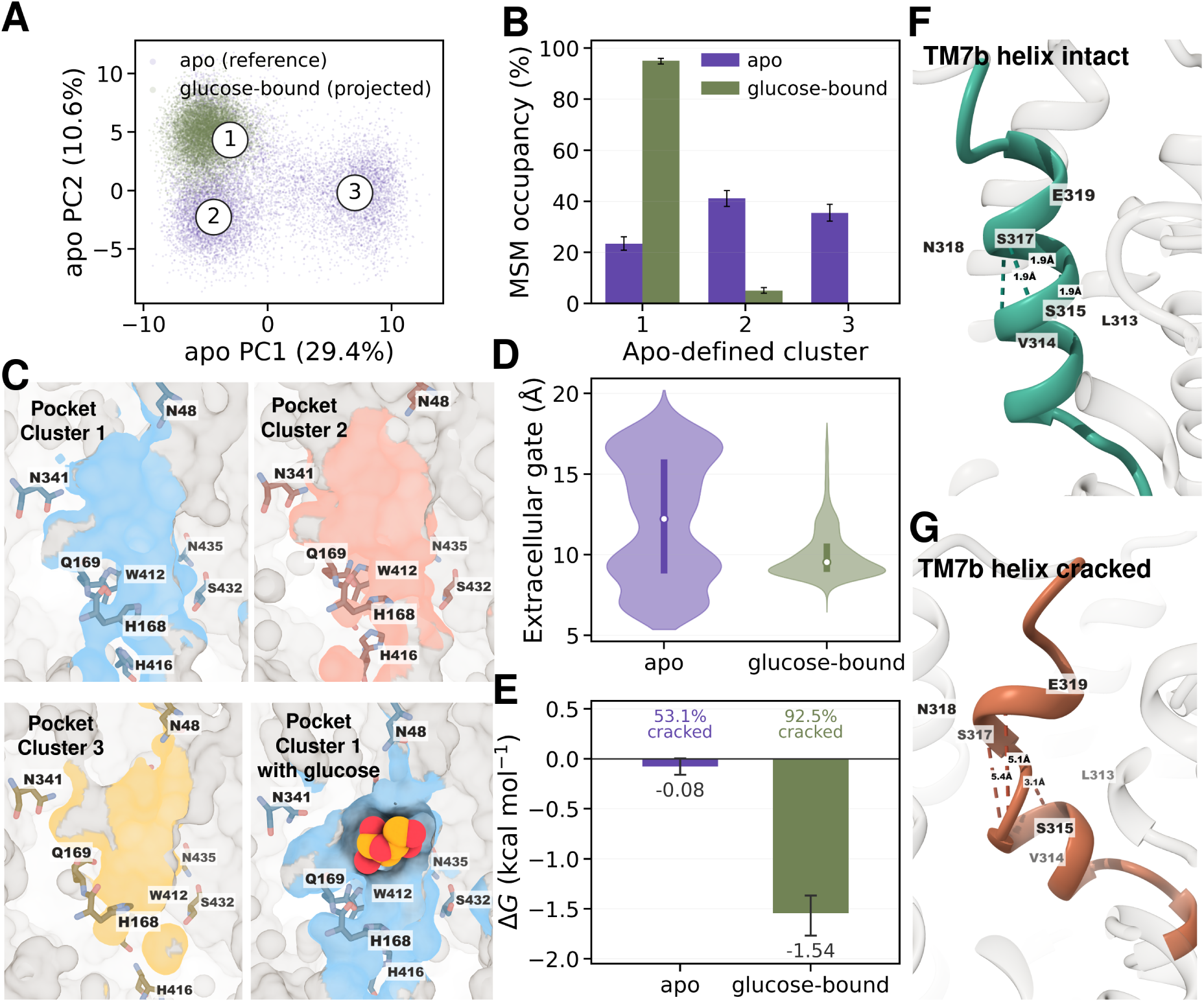
Glucose stabilizes a coupled binding-pocket, gate, and TM7b ensemble in PfHT1. (A) Projection of MSM-weighted apo and glucose-bound pocket conformations onto the first two principal components derived from the apo pocket ensemble. (B) Occupancy of the three apo-defined pocket clusters. (C) Apo structures representing pocket clusters 1–3 and a glucose-bound cluster-1 structure. (D) Distribution of the extracellular-gate distance across all apo and glucose-bound frames. (E) Cracked TM7b populations and equilibrium free-energy differences, Δ*G*_cracked*−*intact_. (F) Intact apo and (G) cracked glucose-bound TM7b structures showing the central backbone *i, i* + 4 hydrogen-bond network.

This narrowing of pocket geometry was accompanied by stabilization of a more restricted global protein conformation. Apo PfHT1 sampled a broad range of extracellular-gate distances, whereas glucose-bound PfHT1 was predominantly confined to distances of approximately 8–10 Å (Figure 6D). Glucose binding was also associated with a pronounced change in the structural integrity of TM7b. Helix cracking was quantified from disruption of the backbone *i, i* + 4 hydrogen-bond network across TM7b residues L313–E319. TM7b was cracked in 53.1% of apo frames and 92.5% of glucose-bound frames. These populations correspond to Δ*G*_cracked*−*intact_ values of *−*0.08 and *−*1.54 kcal mol*^−^*^1^, respectively, producing a glucose-associated shift of *−*1.47 kcal mol*^−^*^1^ (Figure 6E). Representative intact and cracked structures illustrate formation and disruption of the three central hydrogen bonds (Figure 6F,G). Glucose therefore stabilizes a conformational state characterized by a restricted pocket geometry, a partially closed extracellular gate, and a cracked TM7b helix. Previous structural and functional studies indicated that the conserved central binding site alone does not determine substrate specificity in PfHT1. ^15^ Despite its ability to transport several different sugars, PfHT1 coordinates glucose through a binding site that closely resembles those of more selective GLUT transporters. Instead, specificity appears to arise through coupling between initial substrate recognition and the subsequent conformational response of the extracellular gate. The proposed two-step mechanism suggests that initial sugar recognition stabilizes a partially occluded state, after which the substrate promotes an induced-fit transition that completes the binding pocket and enables transport. ^15,17^

Our ensemble analysis provides a dynamic description of this substrate-mediated formation of a favorable binding pocket. The glucose-associated pocket geometry is accessible in apo PfHT1, but it represents only one part of a substantially broader conformational ensemble and is not maintained as a dominant preorganized state. The presence of glucose shifts the ensemble strongly toward this geometry while simultaneously selecting its associated extracellular-gate and TM7b configurations. The favorable glucose-binding pocket can therefore be understood as a substrate-stabilized state that is selected and consolidated from the apo ensemble (Figure 6C).

This mechanism provides a potential explanation for the polyspecificity of PfHT1. The broad apo ensemble offers multiple accessible pocket and gating configurations that can serve as starting points for the recognition of chemically distinct substrates. Different transported sugars may select and stabilize the conformations most compatible with their own stereo-chemistry, effectively helping to form their own transport-competent binding pockets. Pocket diversity therefore provides a structural counterpart to the thermodynamic and kinetic flexibility described above, supporting a model in which different sugars stabilize compatible combinations of pocket geometry and gating conformations.

## Discussion

PfHT1 polyspecificity raises the question of how one transporter accommodates different sugars while maintaining productive alternating access. Our results connect this problem to the substrate-dependent organization of conformational energetics, transition kinetics, and binding-pocket geometry. The glucose-resolved simulations establish how these features change together, while the glucose and fructose-dependent mutational phenotypes support the functional relevance of a residue network whose contributions vary with substrate.

The free energy landscapes demonstrate that PfHT1 follows a canonical MFS-type alternating access mechanism, progressing through outward-facing (OF), occluded (OC), and inward-facing (IF) intermediates. However, distinct differences emerge between apo and holo states. In the apo transporter, flexible gating helices enable exploration of a broad conformational space that includes weakly populated hourglass-associated configurations. Although the HG region does not form a stable equilibrium minimum, iGate opening first configurations can nevertheless contribute to the reactive ensemble as short-lived kinetic corridors. This broad apo ensemble may be particularly relevant to the promiscuous substrate profile of PfHT1, because access to multiple conformational substates provides a structural landscape within which chemically distinct substrates can be accommodated. Substrate binding can then selectively stabilize subsets of this pre-existing ensemble, coupling recognition to productive conformational progression. This conformational breadth extends to the binding pocket: apo PfHT1 populates three distinct pocket clusters, whereas 95% of glucose-bound conformations occupy a single cluster. The accompanying restriction of extracellular-gate distances and enrichment of TM7b cracking indicate that glucose stabilizes a coupled pocket–gate configuration from the broader apo ensemble.

TM7b helix cracking provides a structural mechanism linking local substrate recognition to global conformational change. Its kink, centered around the G309–S315–N316 motif, is strongly associated with extracellular-gate closure and substrate progression. The correlation between TM7b bending and substrate position supports a role for this helix as a mechanical element coupling gating to substrate progression. This finding aligns with recent studies on mammalian GLUT transporters, suggesting that helix flexibility within TM7b is a conserved strategy for lowering energetic barriers in sugar transporters. These observations provide ensemble-level support for the two-step mechanism proposed by Drew and coworkers, in which initial substrate recognition is followed by substrate-mediated stabilization and completion of a transport-competent binding pocket. The glucose-associated geometry is accessible in apo PfHT1 but becomes dominant only after glucose binding.

Kinetic analysis via MSMs quantifies these mechanistic insights. In the apo cycle, multiple routes connect OF and IF states, yet transitions through the canonical OC state are both thermodynamically and kinetically preferred. Glucose binding does not uniformly reduce all MFPTs, instead, it accelerates progression toward IF while slowing key reverse transitions toward OF, thereby converting the mixed directional preference of apo PfHT1 into a rectified glucose-bound transport cycle. These kinetic results provide an atomistic basis for the substrate-dependent bias in conformational exchange.

The transition-path analysis extends the state-based kinetic picture by resolving how reactive current is distributed between OF and IF. Apo PfHT1 accesses two comparably weighted transition tubes with different geometries: one samples nOF-OF before progressing through OC, whereas the other follows a more direct nOF-nOC-IF corridor. Both contain mixtures of eGate-closing-first and iGate-opening-first sequences, showing that geometric pathway heterogeneity cannot be reduced to binary gate-event order. The committor projections confirm that these tubes traverse different conformational regions while progressing between the same endpoints. In glucose-bound PfHT1, both displayed geometric partitions are dominated by eGate closure before iGate opening and converge through the stabilized OC region. Because the unrestricted gap statistic supports one diffuse glucose-bound pathway population, these partitions represent alternative realizations of a common OC-mediated mechanism rather than independent transport mechanisms. Thus, glucose binding reorganizes a heterogeneous apo reactive ensemble into a more strongly funneled conformational pathway.

The integrated residue analysis provides a complementary molecular view of substrate recognition. Direct contact analysis identifies residues that physically interact with glucose as it moves through the transporter, whereas the GAT highlights residue environments that are informative of substrate position and conformational state. The overlap between these analyses identifies positions supported by both direct interaction and broader structural context, while additional GAT-prioritized residues point to distributed contributions beyond the immediate binding pocket. Functional testing further supports the relevance of this residue network, as most alanine-substitution variants reduced PfHT1-dependent growth in glucose, fructose, or both. Importantly, several substitutions produced carbon-source-dependent phenotypes, indicating that individual residues can contribute differently to utilization of distinct substrates. Together, these findings support a model in which PfHT1 polyspecificity depends on the coupling of pocket plasticity, conformational energetics, and transport kinetics. The apo ensemble supplies accessible structural alternatives, while substrate interactions stabilize configurations and pathways compatible with productive transport. Glucose provides a resolved example of this coupling, and the substrate-dependent mutational phenotypes support its relevance to the utilization of different sugars. This framework explains why static binding-site structures alone cannot fully describe polyspecificity and motivates testing how other substrates reshape the same transporter landscape. For antimalarial drug discovery, it shifts attention toward the interactions that connect sugar recognition to productive gating. Ligands that disrupt this coupling or stabilize nonproductive conformations could inhibit nutrient uptake even when the central pocket accommodates multiple sugars, making the machinery that enables substrate versatility a potential therapeutic vulnerability.

## Methods

### Molecular dynamics system setup

The starting structure for PfHT1 simulations was obtained from the Protein Data Bank (PDB ID: 6RW3).^15^ Apo (ligand-free) systems were prepared using CHARMM-GUI, following standard membrane protein protocols.^50^ Missing side chains and unresolved loop regions were modeled using the GalaxyLoop web server.^51^ The glucose ligand was parameterized with the CGenFF force field, and the system was solvated with the TIP3P water model and neutralized with 0.15 M NaCl.^52,53^ The CHARMM36m force field was used to model all atomic interactions.^54^ The transporter was embedded in an asymmetric Plasmodium lipid bilayer that reflects the native membrane composition (Table S2). The *Plasmodium* membrane composition was obtained from lipidometric studies on *Plasmodium berghei*, which is a close evolutionary relative of *Plasmodium falciparum*.^55,56^ The final system box measured approximately [87 × 87 × 133 Å*], containing 103k atoms in total for apo system.

Protonation states of ionizable residues were assigned using PDB2PQR/PROPKA at pH 7. ^57^ Representative frames from the outward-facing (OF) apo simulations were used to generate initial configurations for the holo systems. For holo simulation setup, a glucose molecule was inserted in the water box using Packmol and subsequently parametrized using CHARMM-GUI.^50,58^ Hydrogen mass repartitioning was applied to enable longer 4fs timesteps.^59^ The Plasmodium lipid bilayer was mixed and equilibrated using the VMD Membrane Mixer utility to create 10 replicates.^60^

### Simulation protocol

Energy minimization was performed using 5,000 steps of steepest descent followed by 45,000 steps of conjugate gradient. The systems were gradually heated from 0 K to 310 K over 300,000 steps in the NVT ensemble, followed by 3 ns of equilibration at 310 K under NPT conditions. Positional restraints of 5 kcal mol −1 Å-2 were applied to heavy atoms during the minimization and heating stages. Subsequent equilibration consisted of 50 ns of restrained dynamics and 50 ns of unrestrained dynamics to relax the systems prior to production.

Production simulations were performed using OpenMM 7.7.^61^ A Langevin thermostat and Monte Carlo membrane barostat were used to maintain temperature (310 K) and pressure (1 bar), respectively.^62,63^ Bonds involving hydrogen were constrained with SHAKE, allowing a 4 fs integration timestep.^64^ Long-range electrostatics were computed using Particle Mesh Ewald (PME), with a 12 Å cutoff for short-range interactions.^65^

### Adaptive sampling

To accelerate exploration of the extensive conformational landscape of PfHT1, we employed a multi-strategy adaptive sampling^66^ workflow combining Multi-Agent REAP (MA-REAP), MaxEnt VAMPnet, and Least-Counts sampling schemes.^67–69^ Initial unbiased simulations were used to seed the ensemble for both apo and holo systems. In the initial adaptive rounds, we applied MA-REAP, an exploration-driven reinforcement learning approach, to efficiently identify diverse conformational states associated with large-scale gating and helix rearrangements.^67^ MA-REAP adaptively prioritized starting structures based on novelty and progress metrics, thereby broadening sampling coverage during the early exploration phase. Subsequently, we employed the Maximum Entropy VAMPnet (MaxEnt-VAMPnet) approach to guide sampling toward kinetically relevant regions of the conformational space. ^68^ The VAMPnet model was trained on structural descriptors such as inter-helical distances, bending angles, and gating residues, enabling identification of slow collective motions and transition pathways. MaxEnt weighting ensured that both frequently and rarely visited states contributed proportionally to the expansion of the ensemble. In the final rounds, a Least-Counts selection strategy was implemented to ensure complete coverage and eliminate undersampled regions. Starting conformations were chosen from clusters with the lowest representation in the ensemble, ensuring systematic refinement of the free-energy landscape. This combined adaptive framework enabled efficient coverage of both outward- and inward-facing transitions, capturing intermediate occluded conformations for both apo and holo states. The number of adaptive rounds and total aggregate simulation times in each round are summarized in Table S1. The progress of sampling per round is shown in Figures S10 and S11.

### Markov state model construction

Markov State Models (MSMs) were constructed using DEEPTIME (version 0.4.4) to characterize the conformational dynamics of PfHT1 across both apo and holo ensembles. ^70^ The complete set of structural and dynamic features used for MSM construction is listed in Supplementary Table S3. An iterative, data-driven workflow was used to ensure that the resulting models captured kinetically meaningful transitions while maintaining the Markovian property. Initial MSMs were built from an exhaustive set of structural descriptors, including inter-helical distances, gating distances, and local TM7b bending angles (see Supplementary Table S3). Time-lagged Independent Component Analysis (tICA) was applied to reduce dimensionality and identify the slow collective motions driving conformational transitions. Grid searches were performed over a range of cluster counts, tIC dimensions, and lag times. The VAMP2 score was used as an objective metric to identify the optimal hyperparameter set (Figure S13). The working lag time was chosen as the shortest interval at which implied timescales converged across the dominant slow processes. The resulting tICA free-energy landscape was inspected to ensure that all kinetically relevant states were well-connected, indicating reversible sampling. If disconnected regions were identified, additional trajectories from the adaptive sampling rounds were incorporated until detailed balance was satisfied. Cluster populations from raw simulation data were compared to MSM-reweighted populations to confirm preservation of the underlying phase space. Once convergence criteria were met, final MSMs were constructed for both apo and holo systems using the optimized lag time of 20 ns and a total of 1700 and 900 clusters, respectively. Bayesian MSM was constructed to obtain errors for the implied timescales for the MSMs. (Figure S12). Chapman Kolmogorov tests were performed for validating the constructed MSMs (Figure S14, S15). The reversibility and detailed balance was assessed by comparing *π_i_P_ij_*= *π_j_P_ji_* being valid for all MSM states i and j, and the differences between *π_i_P_ij_* and *π_j_P_ji_*being negligible (Figure S16 and S17).

Macrostates corresponding to outward-facing (OF), occluded (OC), inward-facing (IF), and intermediate conformations were defined by clustering kinetically similar microstates. State equilibrium probabilities from the MSM were used to reconstruct free energy landscapes of the apo and holo systems. Transition path theory was then applied to quantify fluxes between macrostates and to compute mean first passage times (MFPTs), thereby revealing the relative kinetic accessibility of different transport pathways.

### Transition Path Theory

Transition-path theory (TPT) was applied separately to the previously validated apo and glucose-bound PfHT1 Markov state models.^44,45^ Outward-facing (OF) and inward-facing (IF) endpoint ensembles were defined using the mean extracellular-gate (eGate) and intracellular-gate (iGate) distances of each microstate, following the conformational definitions used in Figure 1. The net OF to IF reactive flux was decomposed into the 20,000 highest-flux pathways, which captured 89.6% and 94.5% of the total reactive flux for apo and glucose-bound PfHT1, respectively (Figure S19). Each pathway was represented by a normalized, inverse-density-corrected 30 *×* 30 occupancy histogram in the first two tICA coordinates. Because separate tICA models were constructed for apo and glucose-bound PfHT1, pathway finger-prints and subsequent VAE models were generated independently for the two systems.^71,72^ One two-dimensional variational autoencoder was trained for each system using an 80:20 group-aware training-validation split, with pathways grouped according to their microstate sequences.^73^ TPT pathway flux was retained as separate metadata and was not supplied to the VAE or included in its training objective. Additional details about the VAE model architecture and training are provided in the supplementary information (Figure S18-S20, Table S4, SI methods). Pathways were partitioned from their two-dimensional VAE encoder means using K-means clustering with *k* = 2, the Lloyd algorithm, and 100 centroid initializations. Clustering was performed independently for apo and glucose-bound PfHT1. Number of clusters ranging from one to six clusters were evaluated using the within-cluster sum of squares, silhouette coefficient, and gap statistic.^74–76^ The two groups were reported generically as Cluster 1 and Cluster 2, with cluster numbers assigned post hoc in decreasing order of summed pathway flux. Cluster flux fractions were calculated by summing the TPT fluxes of the constituent pathways. Gate-event sequences were assigned independently of the latent-space clustering. The first microstate along each pathway at which the eGate distance decreased to *≤* 6.25 Å and the first microstate at which the iGate distance increased to *≥* 6.75 Å were identified. Pathways were classified as eGate-closing-first, iGate-opening-first, or concerted when both events first occurred in the same microstate. Gate-order fractions within each cluster were calculated by summing the fluxes of the corresponding pathways among the 20,000 decomposed paths. Transition tubes were constructed by distributing each pathway’s flux uniformly among its visited microstates and summing these contributions within each latent-space cluster. The exact TPT forward committor was projected within each tube to visualize progress from OF (*q*^+^ = 0) to IF (*q*^+^ = 1).

### Graph attention network analysis

To identify residue networks that coordinate glucose transport, we developed a graph attention network (GAT) using MSM-weighted molecular dynamics snapshots spanning the complete transport cycle. A total of 10,000 frames were sampled according to the equilibrium probabilities of the Markov state model to ensure representative coverage of both highly populated and transient conformational states.

Each simulation frame was represented as a weighted residue interaction graph. Protein residues and the glucose molecule were treated as graph nodes. Residue node features consisted of three-dimensional residue center of mass coordinates translated relative to the protein center of mass, thereby removing global translational motion while preserving the protein geometry. The ligand node contained no positional information (zero-valued feature vector) to prevent direct encoding of the prediction target. Two classes of edges were included: residue-residue edges and residue-ligand edges. Edges were constructed using an 8 Å distance cutoff between C*α* atoms for residue-residue interactions and between residue C*α* atoms and the ligand center of mass for residue-ligand interactions. Each edge was assigned a continuous weight according to an exponential distance-decay function,

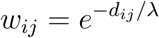

where *d_ij_* is the inter-node distance and *λ*=5 Å. The resulting dataset comprised 10,000 graphs that were randomly divided into training (80%) and test (20%) sets. A graph attention network based on two stacked GATv2 convolutional layers was implemented using PyTorch Geometric.^48,77^ The first layer employed multi-head attention (n=4) followed by a second attention layer to refine node embeddings. Layer normalization and ELU activation functions were applied after each convolutional layer. Graph-level embeddings were obtained using global mean pooling and passed through a two-layer multilayer perceptron (MLP) to predict the ligand z-coordinate, which served as a proxy for substrate position along the transport pathway. Model parameters were optimized using the AdamW optimizer with mean squared error (MSE) loss. Following training, attention coefficients from the second GAT layer were extracted to quantify residue importance throughout the transport cycle. For each graph, attention values associated with residue-to-ligand edges were averaged and grouped according to ligand z-coordinate, generating stage-specific residue importance profiles. Residues exhibiting large changes in attention across substrate positions were interpreted as positions whose local structural environments are informative of substrate progression.

Predicting the ligand position provides a self-supervised learning objective that does not require manually assigned functional labels while forcing the network to learn structural features that distinguish different stages of the transport cycle. Because the ligand node contains no positional information, accurate prediction requires the model to infer substrate location exclusively from the surrounding protein conformation and interaction network. Consequently, the learned attention coefficients therefore provide an interpretable measure of residues contributing strongly to prediction of substrate position at different stages of the conformational cycle.

### Yeast Growth Assay

The pJS288-PfHT1 construct was generated by replacing the ScHXT1-mCherry insert in pJS288 with a yeast codon-optimized sequence encoding Plasmodium falciparum hexose transporter 1 (GenBank: CAA10374.1). The PfHT1 encoding sequence and the pJS288 backbone lacking its insert ScHXT1-mCherry were amplified by polymerase chain reaction (PCR) and assembled using NEBuilder® HiFi DNA Assembly Master Mix (New England Biolabs). The URA3 selection marker in the original plasmid was additionally replaced with HIS3. pJS288 was a gift from Jens Boch (Addgene plasmid # 200714; http://n2t.net/addgene:200714; RRID:Addgene 200714). Single-point mutations were introduced into PfHT1 using overlap-extension PCR and all mutant sequences were verified by plasmid sequencing (Plasmidsaurus). Plasmid-free EBY.VW4000 Saccharomyces cerevisiae cells were maintained in 2% yeast extract, 1% peptone, and 2% maltose (YPM) at 30°C, as previously reported.^78^ PfHT1 wild-type or variant plasmids were transformed into EBY.VW4000 cells following the heat-shock-electroporation (HEEL) method.^79^ Transformants were selected for single colonies on a synthetic defined histidine-free plate [SD(-His); 0.67% yeast nitrogen base with ammonium sulfate and without amino acids,2% agarose; MP Biomedicals] supplemented 1% maltose. Growth assays were performed using clonal transformants in SD(-His) media supplemented with 0.2% glucose or fructose as the test carbon source. Absorbance at 600nm (A600) was measured using a SpectraMax iD5 multi-mode microplate reader (Molecular Devices). Cultures of all mutants, the WT control, and the empty-vector control were diluted to comparable initial cell densities, corresponding to plate reader A600 values of approximately 0.05 to 0.08. The apparent early exponential-phase growth rate was calculated from four biological replicates and used as a proxy for transporter-mediated sugar uptake activity. For comparisons of substrate selectivity among PfHT1 variants, growth rate under each sugar condition was normalized relative to the WT and empty-vector controls, with the WT sugar uptake activity defined as 1, and the empty-vector control activity defined as 0.

### Binding pocket ensemble analysis

Binding-pocket conformations were analysed using 10,000 MSM-weighted apo frames and 10,000 MSM-weighted glucose-bound frames in which glucose remained in the central pocket. Pocket geometry was represented by the 55 pairwise side-chain-centre distances and 55 pairwise C*α* distances among N48, H168, Q169, Q305, Q306, N311, N341, W412, H416, S432, and N435. Features were standardized using the apo ensemble, and PCA and three-cluster MiniBatch K-means models were fitted to the apo frames. Glucose-bound frames were then projected and assigned without refitting. A backbone *i, i* + 4 hydrogen bond was considered formed when the H···O distance was below 2.5 Å and the N–H···O angle exceeded 120*^◦^*. TM7b was classified as cracked when at least two of the three central hydrogen bonds spanning residues 313–319 were absent. Equilibrium free energies were calculated at 310 K as Δ*G*_cracked−intact_ = −*k*_B_*T* ln(*P_cracked_*/*P_intact_*).

### Trajectory analysis

All trajectory analysis were performed using in house python scripts, CPPTRAJ, HOLE package and GetContacts packages.^80–82^ Protein structures were visualized using Pymol 3, ChimeraX and VMD 1.9.4a55.^83–85^

## Data Availability

All simulation data as well as features used for MSM construction are available on request. Code for Graph Attention Model and VAE-based latent path clustering is available here https://github.com/ShuklaGroup/PfHT1. Due to large size of simulation trajectories generated for this study, they will be made available upon request.

## Author Contribution

A.P and D.S. conceived and designed the study. A.P. collected and analyzed the data. L.X. performed yeast growth experiments. A.P. and L.X. drafted the manuscript with help from D.S. D.S. supervised the study and secured funding for the project.

## Acknowledgement

The authors acknowledge support from the National Institutes of Health Award R35GM142745. This manuscript is the result of funding in whole or in part by the National Institutes of Health (NIH). It is subject to the NIH Public Access Policy. Through acceptance of this federal funding, NIH has been given a right to make this manuscript publicly available in PubMed Central upon the Official Date of Publication, as defined by NIH. The authors would like to thank Dr Austin T. Weigle for the helpful discussions and help with membrane composition of plasmodium membrane. The authors also acknowledge support from citizen scientists providing computing hours for simulations performed on Folding@Home, which have enabled us to collect data on a large scale for this project. The authors also thank Prof. Kai Zhang in the Biochemistry Department at the School of Molecular and Cellular Biology at UIUC for providing access to the SpectraMax iD5 plate reader. The authors also thank Prof. Eckhard Boles from the Goethe University, Frankfurt for sharing yeast strain EBY.VW4000.

The authors acknowledge the use of GPT-5.6 (OpenAI) to assist with improving the language, flow, and clarity of the manuscript. The authors reviewed, edited, and approved all AI-assisted text and take full responsibility for the content of this publication.

## Competing Interests

The authors declare that they have no competing interests.

## Supporting Information Available

Details of Markov state models, error plots, simulation details, adaptive sampling details, plasmodium membrane composition, VAE-path clustering models and additional methods.

## Notes

### Competing Interest Statement

The authors have declared no competing interest.

### Summary of Updates

Revised the title and added a section at the end about structural diversity of binding pocket.

## References

(1) Drew, D.; North, R. A.; Nagarathinam, K.; Tanabe, M. Structures and General Transport Mechanisms by the Major Facilitator Superfamily (MFS). Chemical Reviews 2021, 121, 5289–5335.

(2) Beckstein, O.; Naughton, F. General principles of secondary active transporter function. Biophysics Reviews 2022, 3.

(3) Sauve, S.; Williamson, J.; Polasa, A.; Moradi, M. Ins and Outs of Rocker Switch Mechanism in Major Facilitator Superfamily of Transporters. Membranes 2023, 13, 462.

(4) Drew, D.; Boudker, O. Shared Molecular Mechanisms of Membrane Transporters. Annual Review of Biochemistry 2016, 85, 543–572.

(5) Maiden, M. C. J.; Davis, E. O.; Baldwin, S. A.; Moore, D. C. M.; Henderson, P. J. F. Mammalian and bacterial sugar transport proteins are homologous. Nature 1987, 325, 641–643.

(6) Deng, D.; Xu, C.; Sun, P.; Wu, J.; Yan, C.; Hu, M.; Yan, N. Crystal structure of the human glucose transporter GLUT1. Nature 2014, 510, 121–125.

(7) Deng, D.; Sun, P.; Yan, C.; Ke, M.; Jiang, X.; Xiong, L.; Ren, W.; Hirata, K.; Yamamoto, M.; Fan, S.; Yan, N. Molecular basis of ligand recognition and transport by glucose transporters. Nature 2015, 526, 391–396.

(8) Yuan, Y.; Kong, F.; Xu, H.; Zhu, A.; Yan, N.; Yan, C. Cryo-EM structure of human glucose transporter GLUT4. Nature Communications 2022, 13.

(9) Nomura, N. et al. Structure and mechanism of the mammalian fructose transporter GLUT5. Nature 2015, 526, 397–401.

(10) Lee, S. S.; Kim, S.; Jin, M. S. Cryo-EM structure of the human glucose transporter GLUT7. Biochemical and Biophysical Research Communications 2024, 738, 150544.

(11) Shen, Z.; Xu, L.; Wu, T.; Wang, H.; Wang, Q.; Ge, X.; Kong, F.; Huang, G.; Pan, X. Structural basis for urate recognition and apigenin inhibition of human GLUT9. Nature Communications 2024, 15.

(12) Sun, L.; Zeng, X.; Yan, C.; Sun, X.; Gong, X.; Rao, Y.; Yan, N. Crystal structure of a bacterial homologue of glucose transporters GLUT1–4. Nature 2012, 490, 361–366.

(13) Quistgaard, E. M.; Löw, C.; Moberg, P.; Trésaugues, L.; Nordlund, P. Structural basis for substrate transport in the GLUT-homology family of monosaccharide transporters. Nature Structural & Molecular Biology 2013, 20, 766–768.

(14) Wisedchaisri, G.; Park, M.-S.; Iadanza, M. G.; Zheng, H.; Gonen, T. Proton-coupled sugar transport in the prototypical major facilitator superfamily protein XylE. Nature Communications 2014, 5.

(15) Qureshi, A. A.; Suades, A.; Matsuoka, R.; Brock, J.; McComas, S. E.; Nji, E.; Orellana, L.; Claesson, M.; Delemotte, L.; Drew, D. The molecular basis for sugar import in malaria parasites. Nature 2020, 578, 321–325.

(16) Jiang, X. et al. Structural Basis for Blocking Sugar Uptake into the Malaria Parasite Plasmodium falciparum. Cell 2020, 183, 258–268.e12.

(17) Ahn, D.-H.; Alleva, C.; Reichenbach, T.; Gulati, A.; Ruda, A.; Bonaccorsi, M.; Silberberg, J. M.; Claesson, M.; Suades, A.; Delemotte, L.; Widmalm, G.; Drew, D. A two-step mechanism for sugar translocation. Nature Structural & Molecular Biology 2026,

(18) Paulsen, P. A.; Custódio, T. F.; Pedersen, B. P. Crystal structure of the plant symporter STP10 illuminates sugar uptake mechanism in monosaccharide transporter superfamily. Nature Communications 2019, 10.

(19) Bavnhøj, L.; Paulsen, P. A.; Flores-Canales, J. C.; Schiøtt, B.; Pedersen, B. P. Molecular mechanism of sugar transport in plants unveiled by structures of glucose/H+ symporter STP10. Nature Plants 2021, 7, 1409–1419.

(20) Andersen, C. G.; Bavnhøj, L.; Brag, S.; Bohush, A.; Chrenková, A.; Driller, J. H.; Pedersen, B. P. Comparative analysis of STP6 and STP10 unravels molecular selectivity in sugar transport proteins. Proceedings of the National Academy of Sciences 2025, 122.

(21) Bavnhøj, L.; Driller, J. H.; Zuzic, L.; Stange, A. D.; Schiøtt, B.; Pedersen, B. P. Structure and sucrose binding mechanism of the plant SUC1 sucrose transporter. Nature Plants 2023, 9, 938–950.

(22) Yan, N. A Glimpse of Membrane Transport through Structures—Advances in the Structural Biology of the GLUT Glucose Transporters. Journal of Molecular Biology 2017, 429, 2710–2725.

(23) Kazmier, K.; Claxton, D. P.; Mchaourab, H. S. Alternating access mechanisms of LeuT-fold transporters: trailblazing towards the promised energy landscapes. Current Opinion in Structural Biology 2017, 45, 100–108.

(24) Claxton, D. P.; Jagessar, K. L.; Mchaourab, H. S. Principles of Alternating Access in Multidrug and Toxin Extrusion (MATE) Transporters. Journal of Molecular Biology 2021, 433, 166959.

(25) Mitrovic, D.; McComas, S. E.; Alleva, C.; Bonaccorsi, M.; Drew, D.; Delemotte, L. Reconstructing the transport cycle in the sugar porter superfamily using coevolution-powered machine learning. eLife 2023, 12.

(26) McComas, S. E.; Reichenbach, T.; Mitrovic, D.; Alleva, C.; Bonaccorsi, M.; Delemotte, L.; Drew, D. Determinants of sugar-induced influx in the mammalian fructose transporter GLUT5. eLife 2023, 12.

(27) Woodrow, C. J.; Penny, J. I.; Krishna, S. Intraerythrocytic Plasmodium falciparum Expresses a High Affinity Facilitative Hexose Transporter. Journal of Biological Chemistry 1999, 274, 7272–7277.

(28) Woodrow, C. J.; Burchmore, R. J.; Krishna, S. Hexose permeation pathways in Plasmodium falciparum - infected erythrocytes. Proceedings of the National Academy of Sciences 2000, 97, 9931–9936.

(29) Kirk, K. Glucose uptake in Plasmodium falciparum-infected erythrocytes is an equilibrative not an active process. Molecular and Biochemical Parasitology 1996, 82, 195–205.

(30) Miyashita, O.; Onuchic, J. N.; Wolynes, P. G. Nonlinear elasticity, proteinquakes, and the energy landscapes of functional transitions in proteins. Proceedings of the National Academy of Sciences 2003, 100, 12570–12575.

(31) Whitford, P. C.; Miyashita, O.; Levy, Y.; Onuchic, J. N. Conformational Transitions of Adenylate Kinase: Switching by Cracking. Journal of Molecular Biology 2007, 366, 1661–1671.

(32) Joët, T.; Eckstein-Ludwig, U.; Morin, C.; Krishna, S. Validation of the hexose transporter of Plasmodium falciparum as a novel drug target. Proceedings of the National Academy of Sciences 2003, 100, 7476–7479.

(33) Blume, M.; Hliscs, M.; Rodriguez-Contreras, D.; Sanchez, M.; Landfear, S.; Lucius, R.; Matuschewski, K.; Gupta, N. A constitutive pan-hexose permease for the Plasmodium life cycle and transgenic models for screening of antimalarial sugar analogs. The FASEB Journal 2010, 25, 1218–1229.

(34) Chan, M. C.; Shukla, D. Markov state modeling of membrane transport proteins. Journal of Structural Biology 2021, 213, 107800.

(35) Shukla, D.; Hernández, C. X.; Weber, J. K.; Pande, V. S. Markov State Models Provide Insights into Dynamic Modulation of Protein Function. Accounts of Chemical Research 2015, 48, 414–422.

(36) Husic, B. E.; Pande, V. S. Markov State Models: From an Art to a Science. Journal of the American Chemical Society 2018, 140, 2386–2396.

(37) Konovalov, K. A.; Unarta, I. C.; Cao, S.; Goonetilleke, E. C.; Huang, X. Markov State Models to Study the Functional Dynamics of Proteins in the Wake of Machine Learning. JACS Au 2021, 1, 1330–1341.

(38) Dean, T. J.; Feng, J.; Shukla, D. How Minor Sequence Changes Enable Mechanistic Diversity in MFS Transporters? An Atomic-Level Rationale for Symport Emergence in NarU. Journal of Chemical Information and Modeling 2026, 66, 2358–2368.

(39) Razavi, A. M.; Khelashvili, G.; Weinstein, H. A Markov State-based Quantitative Kinetic Model of Sodium Release from the Dopamine Transporter. Scientific Reports 2017, 7.

(40) Chan, M. C.; Alfawaz, Y.; Paul, A.; Shukla, D. Molecular insights into the elevator-type mechanism of the cyanobacterial bicarbonate transporter BicA. Biophysical Journal 2025, 124, 379–392.

(41) Cheng, K. J.; Selvam, B.; Chen, L.-Q.; Shukla, D. Distinct Substrate Transport Mechanism Identified in Homologous Sugar Transporters. The Journal of Physical Chemistry B 2019, 123, 8411–8418.

(42) Weigle, A. T.; Shukla, D. The Arabidopsis AtSWEET13 transporter discriminates sugars by selective facial and positional substrate recognition. Communications Biology 2024, 7.

(43) Feng, J.; Selvam, B.; Shukla, D. How do antiporters exchange substrates across the cell membrane? An atomic-level description of the complete exchange cycle in NarK. Structure 2021, 29, 922–933.e3.

(44) Metzner, P.; Schütte, C.; Vanden-Eijnden, E. Transition Path Theory for Markov Jump Processes. Multiscale Modeling & Simulation 2009, 7, 1192–1219.

(45) Noé, F.; Schütte, C.; Vanden-Eijnden, E.; Reich, L.; Weikl, T. R. Constructing the equilibrium ensemble of folding pathways from short off-equilibrium simulations. Proceedings of the National Academy of Sciences 2009, 106, 19011–19016.

(46) Pindi, C.; Ahsan, M.; Sinha, S.; Palermo, G. Graph Attention Neural Networks Reveal TnsC Filament Assembly in a CRISPR-Associated Transposon. 2025,

(47) Ahsan, M.; Pindi, C.; Sinha, S.; Patel, A. C.; Palermo, G. Graph neural networks for molecular dynamics simulations. Current Opinion in Structural Biology 2026, 97, 103238.

(48) Veličković, P.; Cucurull, G.; Casanova, A.; Romero, A.; Liò, P.; Bengio, Y. Graph Attention Networks. 2017; https://arxiv.org/abs/1710.10903.

(49) Huang, J. et al. Orthosteric–allosteric dual inhibitors of PfHT1 as selective antimalarial agents. Proceedings of the National Academy of Sciences 2021, 118.

(50) Jo, S.; Kim, T.; Iyer, V. G.; Im, W. CHARMM-GUI: A web-based graphical user interface for CHARMM. Journal of Computational Chemistry 2008, 29, 1859–1865.

(51) Ko, J.; Park, H.; Heo, L.; Seok, C. GalaxyWEB server for protein structure prediction and refinement. Nucleic Acids Research 2012, 40, W294–W297.

(52) Vanommeslaeghe, K.; Hatcher, E.; Acharya, C.; Kundu, S.; Zhong, S.; Shim, J.; Darian, E.; Guvench, O.; Lopes, P.; Vorobyov, I.; Mackerell, A. D. CHARMM general force field: A force field for drug-like molecules compatible with the CHARMM all-atom additive biological force fields. Journal of Computational Chemistry 2009, 31, 671–690.

(53) Jorgensen, W. L.; Chandrasekhar, J.; Madura, J. D.; Impey, R. W.; Klein, M. L. Comparison of simple potential functions for simulating liquid water. The Journal of Chemical Physics 1983, 79, 926–935.

(54) Huang, J.; Rauscher, S.; Nawrocki, G.; Ran, T.; Feig, M.; de Groot, B. L.; Grubmüller, H.; MacKerell, A. D. CHARMM36m: an improved force field for folded and intrinsically disordered proteins. Nature Methods 2016, 14, 71–73.

(55) Wallace, W. R.; Finerty, J. F.; Dimopoullos, G. T. Studies on the Lipids of Plasmodium Lophurae and Plasmodium Berghei. The American Journal of Tropical Medicine and Hygiene 1965, 14, 715–718.

(56) Wallace, W. R. Fatty Acid Composition of Lipid Classes in Plasmodium Lophurae and Plasmodium Berghei. The American Journal of Tropical Medicine and Hygiene 1966, 15, 811–813.

(57) Olsson, M. H. M.; Søndergaard, C. R.; Rostkowski, M.; Jensen, J. H. PROPKA3: Consistent Treatment of Internal and Surface Residues in Empirical pKa Predictions. Journal of Chemical Theory and Computation 2011, 7, 525–537.

(58) Martínez, L.; Andrade, R.; Birgin, E. G.; Martínez, J. M. PACKMOL: A package for building initial configurations for molecular dynamics simulations. Journal of Computational Chemistry 2009, 30, 2157–2164.

(59) Hopkins, C. W.; Le Grand, S.; Walker, R. C.; Roitberg, A. E. Long-Time-Step Molecular Dynamics through Hydrogen Mass Repartitioning. Journal of Chemical Theory and Computation 2015, 11, 1864–1874.

(60) Licari, G.; Dehghani-Ghahnaviyeh, S.; Tajkhorshid, E. Membrane Mixer: A Toolkit for Efficient Shuffling of Lipids in Heterogeneous Biological Membranes. Journal of Chemical Information and Modeling 2022, 62, 986–996.

(61) Eastman, P.; Swails, J.; Chodera, J. D.; McGibbon, R. T.; Zhao, Y.; Beauchamp, K. A.; Wang, L.-P.; Simmonett, A. C.; Harrigan, M. P.; Stern, C. D.; Wiewiora, R. P.; Brooks, B. R.; Pande, V. S. OpenMM 7: Rapid development of high performance algorithms for molecular dynamics. PLOS Computational Biology 2017, 13, e1005659.

(62) Davidchack, R. L.; Handel, R.; Tretyakov, M. V. Langevin thermostat for rigid body dynamics. The Journal of Chemical Physics 2009, 130.

(63) Åqvist, J.; Wennerström, P.; Nervall, M.; Bjelic, S.; Brandsdal, B. O. Molecular dynamics simulations of water and biomolecules with a Monte Carlo constant pressure algorithm. Chemical Physics Letters 2004, 384, 288–294.

(64) Ryckaert, J.-P.; Ciccotti, G.; Berendsen, H. J. Numerical integration of the cartesian equations of motion of a system with constraints: molecular dynamics of n-alkanes. Journal of Computational Physics 1977, 23, 327–341.

(65) Darden, T.; York, D.; Pedersen, L. Particle mesh Ewald: An Nlog(N) method for Ewald sums in large systems. The Journal of Chemical Physics 1993, 98, 10089–10092.

(66) Kleiman, D. E.; Nadeem, H.; Shukla, D. Adaptive Sampling Methods for Molecular Dynamics in the Era of Machine Learning. The Journal of Physical Chemistry B 2023, 127, 10669–10681.

(67) Kleiman, D. E.; Shukla, D. Multiagent Reinforcement Learning-Based Adaptive Sampling for Conformational Dynamics of Proteins. Journal of Chemical Theory and Computation 2022, 18, 5422–5434.

(68) Kleiman, D. E.; Shukla, D. Active Learning of the Conformational Ensemble of Proteins Using Maximum Entropy VAMPNets. Journal of Chemical Theory and Computation 2023, 19, 4377–4388.

(69) Nadeem, H.; Shukla, D. Ensemble Adaptive Sampling Scheme: Identifying an Optimal Sampling Strategy via Policy Ranking. Journal of Chemical Theory and Computation 2025, 21, 4626–4639.

(70) Hoffmann, M.; Scherer, M.; Hempel, T.; Mardt, A.; de Silva, B.; Husic, B. E.; Klus, S.; Wu, H.; Kutz, N.; Brunton, S. L.; Noé, F. Deeptime: a Python library for machine learning dynamical models from time series data. Machine Learning: Science and Technology 2021, 3, 015009.

(71) Qiu, Y.; O’Connor, M. S.; Xue, M.; Liu, B.; Huang, X. An Efficient Path Classification Algorithm Based on Variational Autoencoder to Identify Metastable Path Channels for Complex Conformational Changes. Journal of Chemical Theory and Computation 2023, 19, 4728–4742.

(72) Yin, S.; Mi, X.; Barrett, S. E.; Mitchell, D. A.; Shukla, D. De novo Folding Mechanisms of Lasso Peptides. *bioRxiv* 2026, doi: 10.64898/2026.03.30.715466.

(73) Kingma, D. P.; Welling, M. Auto-Encoding Variational Bayes. 2013; https://arxiv.org/abs/1312.6114.

(74) Lloyd, S. Least squares quantization in PCM. IEEE Transactions on Information Theory 1982, 28, 129–137.

(75) Rousseeuw, P. J. Silhouettes: A graphical aid to the interpretation and validation of cluster analysis. Journal of Computational and Applied Mathematics 1987, 20, 53–65.

(76) Tibshirani, R.; Walther, G.; Hastie, T. Estimating the Number of Clusters in a Data Set Via the Gap Statistic. Journal of the Royal Statistical Society Series B: Statistical Methodology 2001, 63, 411–423.

(77) Fey, M.; Lenssen, J. E. Fast Graph Representation Learning with PyTorch Geometric. 2019; https://arxiv.org/abs/1903.02428.

(78) Wieczorke, R.; Krampe, S.; Weierstall, T.; Freidel, K.; Hollenberg, C. P.; Boles, E. Concurrent knock-out of at least 20 transporter genes is required to block uptake of hexoses in ¡i¿Saccharomyces cerevisiae¡/i¿. FEBS Letters 1999, 464, 123–128.

(79) Wäneskog, M.; Hoch-Schneider, E. E.; Garg, S.; Kronborg Cantalapiedra, C.; Schäfer, E.; Krogh Jensen, M.; Damgaard Jensen, E. Accurate phenotype-to-genotype mapping of high-diversity yeast libraries by heat-shock-electroporation (HEEL). mBio 2025, 16.

(80) Roe, D. R.; Cheatham, T. E. PTRAJ and CPPTRAJ: Software for Processing and Analysis of Molecular Dynamics Trajectory Data. Journal of Chemical Theory and Computation 2013, 9, 3084–3095.

(81) Smart, O. S.; Neduvelil, J. G.; Wang, X.; Wallace, B.; Sansom, M. S. HOLE: A program for the analysis of the pore dimensions of ion channel structural models. Journal of Molecular Graphics 1996, 14, 354–360.

(82) GetContacts package. https://getcontacts.github.io/.

(83) Schrödinger, L. PyMOL. http://www.pymol.org/pymol.

(84) Meng, E. C.; Goddard, T. D.; Pettersen, E. F.; Couch, G. S.; Pearson, Z. J.; Morris, J. H.; Ferrin, T. E. UCSF ChimeraX: Tools for structure building and analysis. Protein Science 2023, 32.

(85) Humphrey, W.; Dalke, A.; Schulten, K. VMD – Visual Molecular Dynamics. Journal of Molecular Graphics 1996, 14, 33–38.

