## supporting information for "Thermodynamic, Kinetic, and Structural Basis of Polyspecificity in the Malaria Sugar Transporter PfHT1"

Arnav Paul,[†] Lingyun Xu,[‡] and Diwakar Shukla[*,¶,†,§,‡]

†*Department of Chemistry, University of Illinois at Urbana-Champaign, Urbana, Illinois 61801, United States*

‡*Department of Bioengineering, University of Illinois Urbana-Champaign, Urbana, IL 61801, United States*

¶*Department of Chemical and Biomolecular Engineering, University of Illinois at Urbana-Champaign, Champaign, Illinois 61801, United States*

§*Department of Plant Biology, University of Illinois at Urbana-Champaign, Champaign, Illinois 61801, United States*

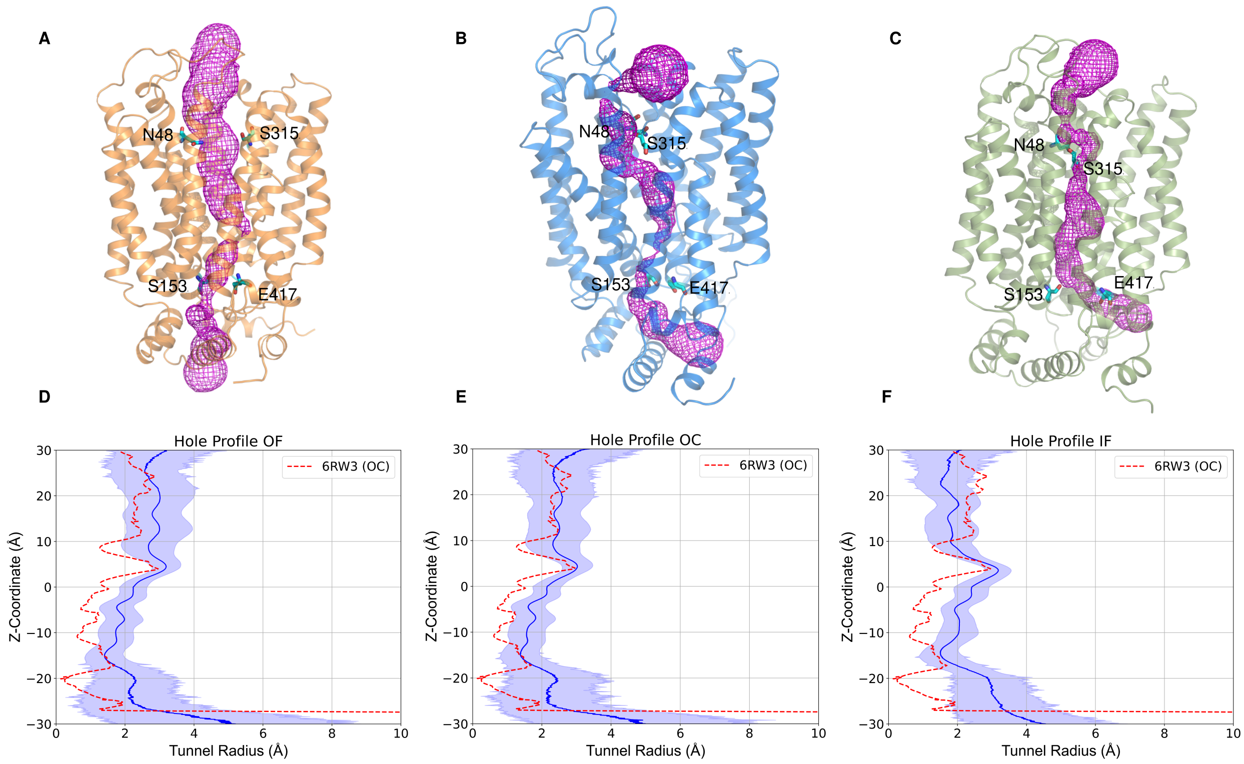


Figure 1: Hole profile: Representative structures showing the accessible volume in the transport tunnel and gating residues for (A) OF state, (B) OC state and (C) IF state. Hole profile for ensemble of structures compared to the resolved structure (PDB: 6RW3) for (D) OF, (E) OC and (F) IF states.

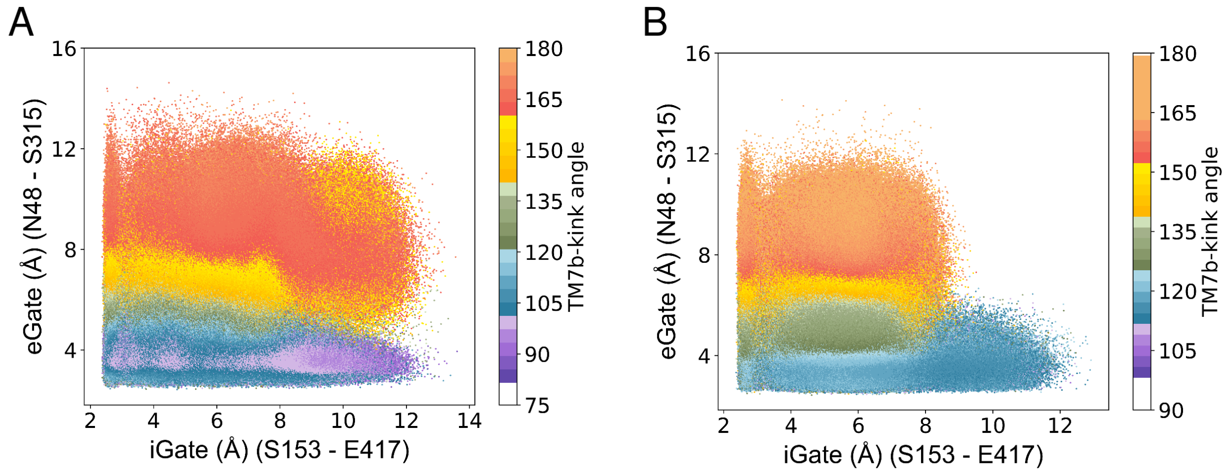


Figure 2: TM7b kink angle projected on the gating landscape highlighting correlation of eGate with TM7b kink angle for (A) apo simulations and (B) holo simulations.

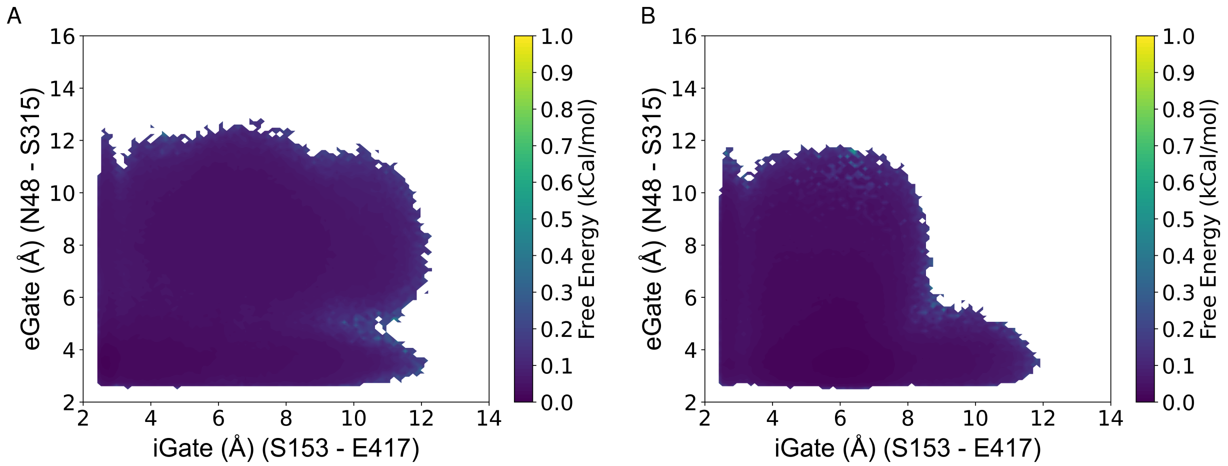


Figure 3: Error plots for the gating free energy landscape by bootstrapping 80% of the total simulation data 500 times for A) Apo cycle and B)Holo cycle.

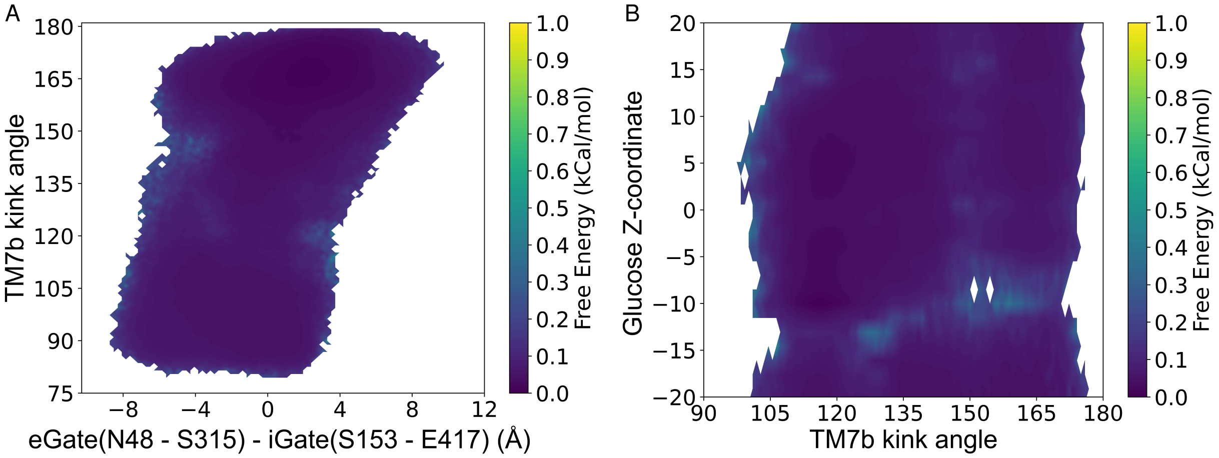


Figure 4: Error plots for the free energy landscape by bootstrapping 80% of the total simulation data 500 times for A) the correlation of TM7b with the conformational change of PfHT1 and (B) Free energy landscape of glucose transport projected on the TM7b kink angle.

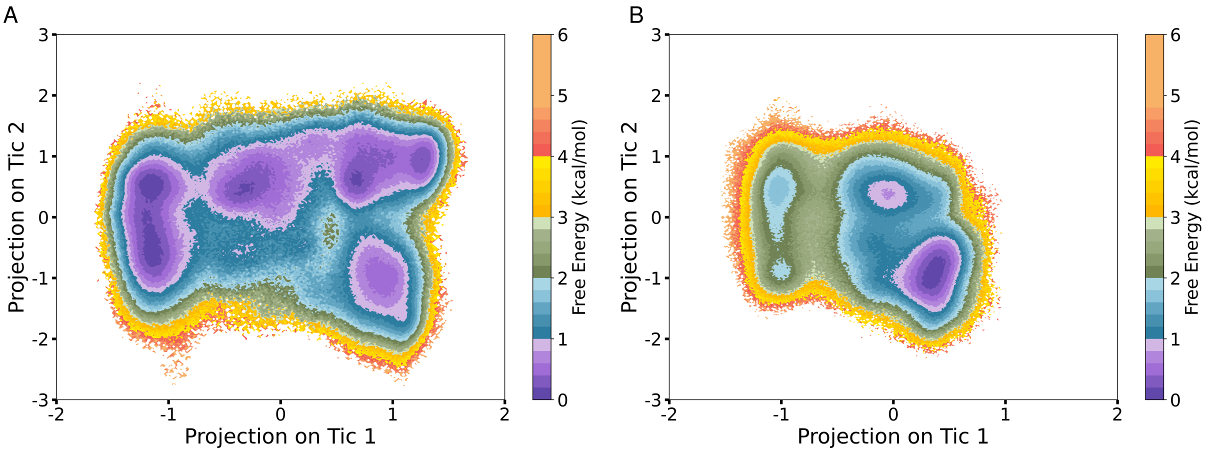


Figure 5: TICA (time-lagged independent component analysis) plot for PfHT1 conformational cycle projected on the first two time-lagged independent components (tICs). A) The tIC projection of apo simulation data on tic1 and tic2 and B) holo data projected on apo tic1 and tic2.

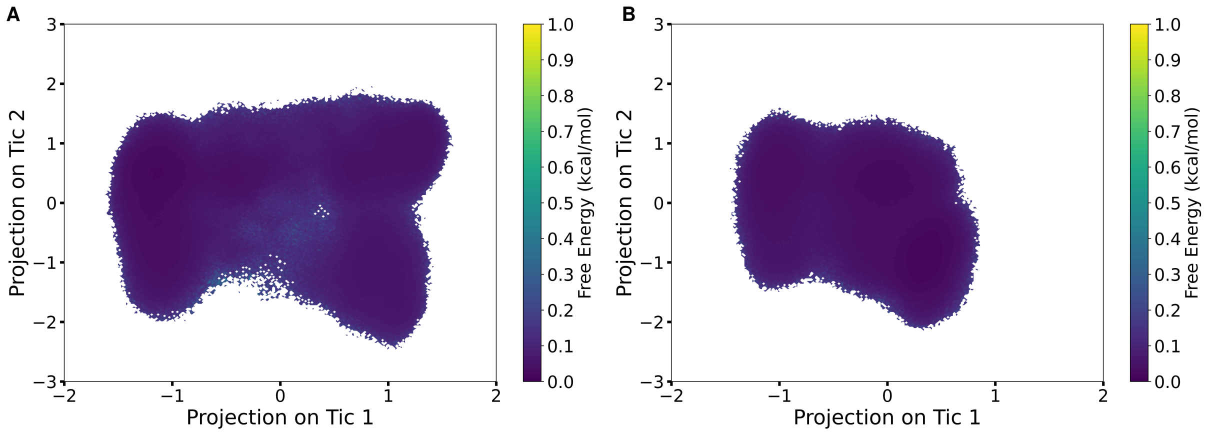


Figure 6: Error plots for the gating free energy landscape by bootstrapping 80% of the total simulation data 500 times for A) The tIC projection of apo simulation data on tic1 and tic2 and B) tic projection of holo simulation data projected on apo tic1 and tic2.

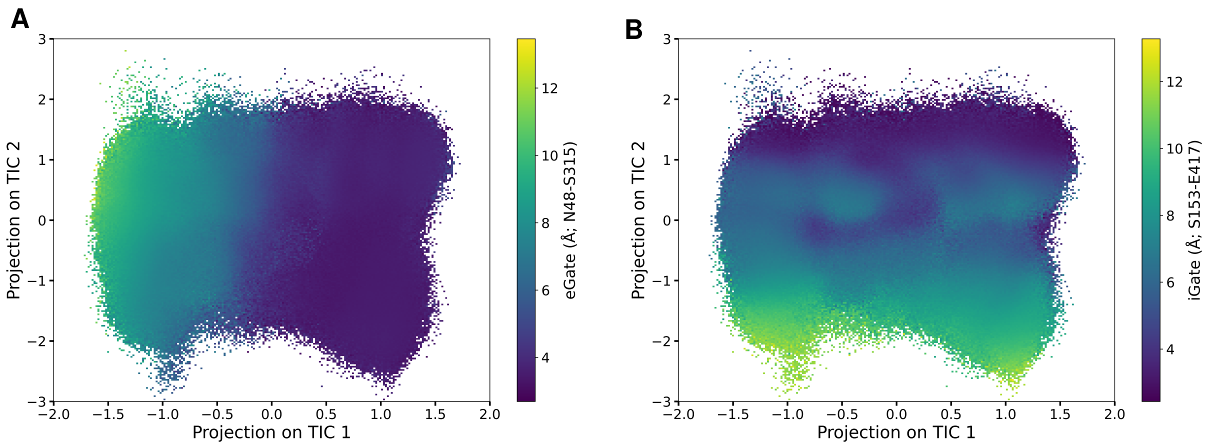


Figure 7: Correlation of gating distances with the tIC dimensions for apo simulation data. (A) Extracellular gating distances projected on the tic plots shows strong correlation with extracellular gating distances with tic1. (B) Intracellular gating distances projected on the tic plots shows strong correlation with intracellular gating distances with tic2.

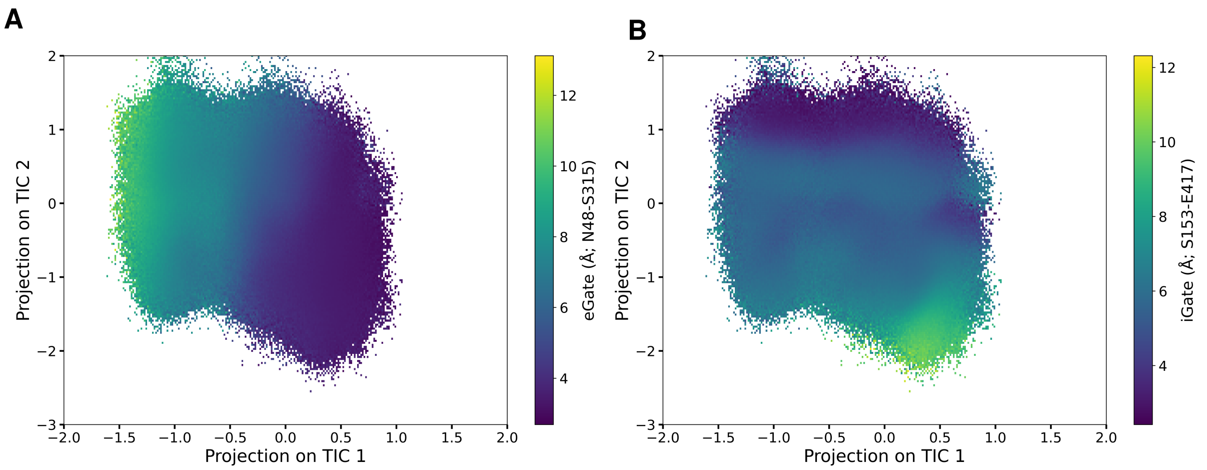


Figure 8: Correlation of gating distances with the tIC dimensions for holo simulation data. (A) Extracellular gating distances projected on the tic plots shows strong correlation with extracellular gating distances with tic1. (B) Intracellular gating distances projected on the tic plots shows strong correlation with intracellular gating distances with tic2.

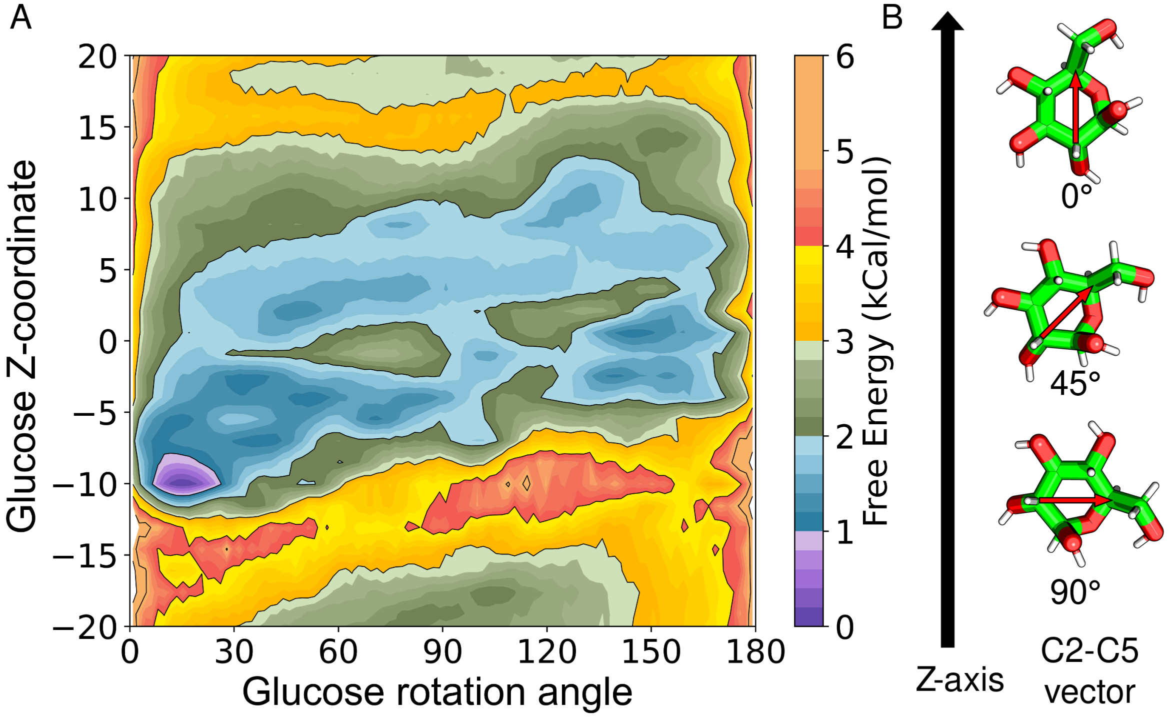


Figure 9: (A) Free energy landscape for glucose z-coordinate vs glucose rotation angle describing the preferred orientation of glucose throughout the transport process and (B) Glucose rotation angle calculated by calculating the angle between the C2-C5 atom vector of glucose the Z-axis.

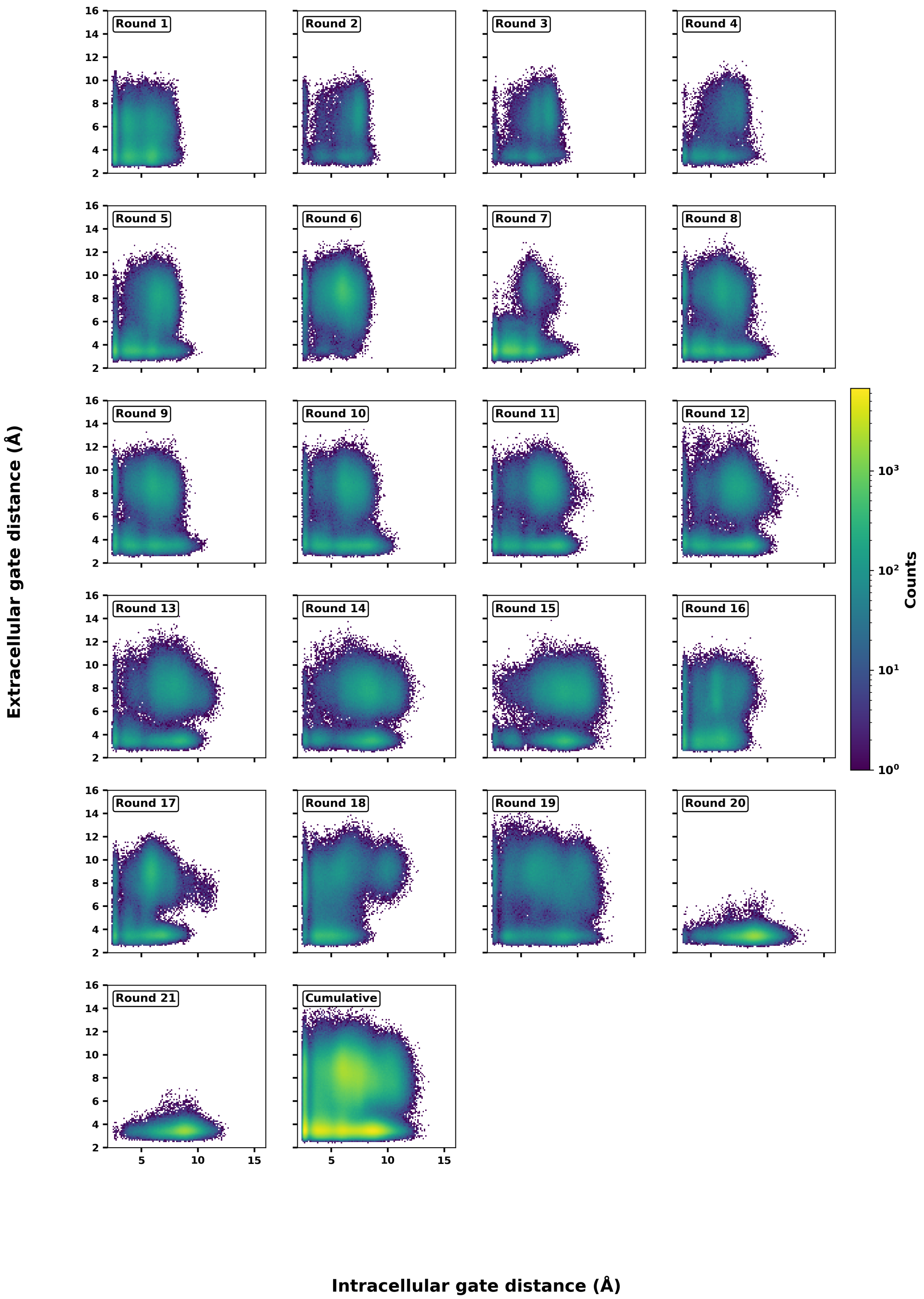


Figure 10: Progression of adaptive sampling per round projected on the eGate and iGate distances. Total of 21 rounds of adaptive sampling totaling 380$\mu$s of simulation for the apo system.

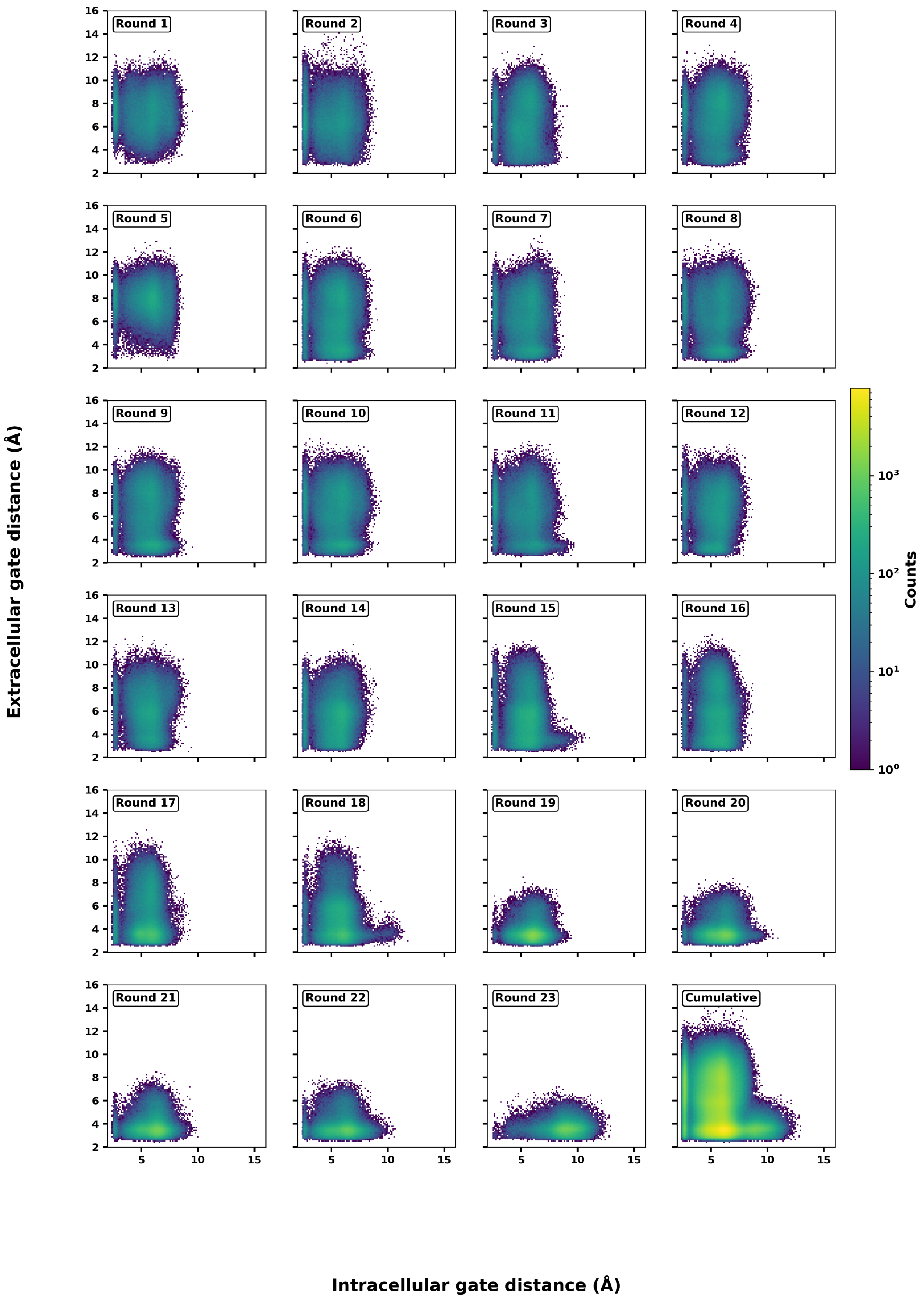


Figure 11: Progression of adaptive sampling per round projected on the eGate and iGate distances. Total of 23 rounds of adaptive sampling totaling 460$\mu$s of simulation for the holo system.

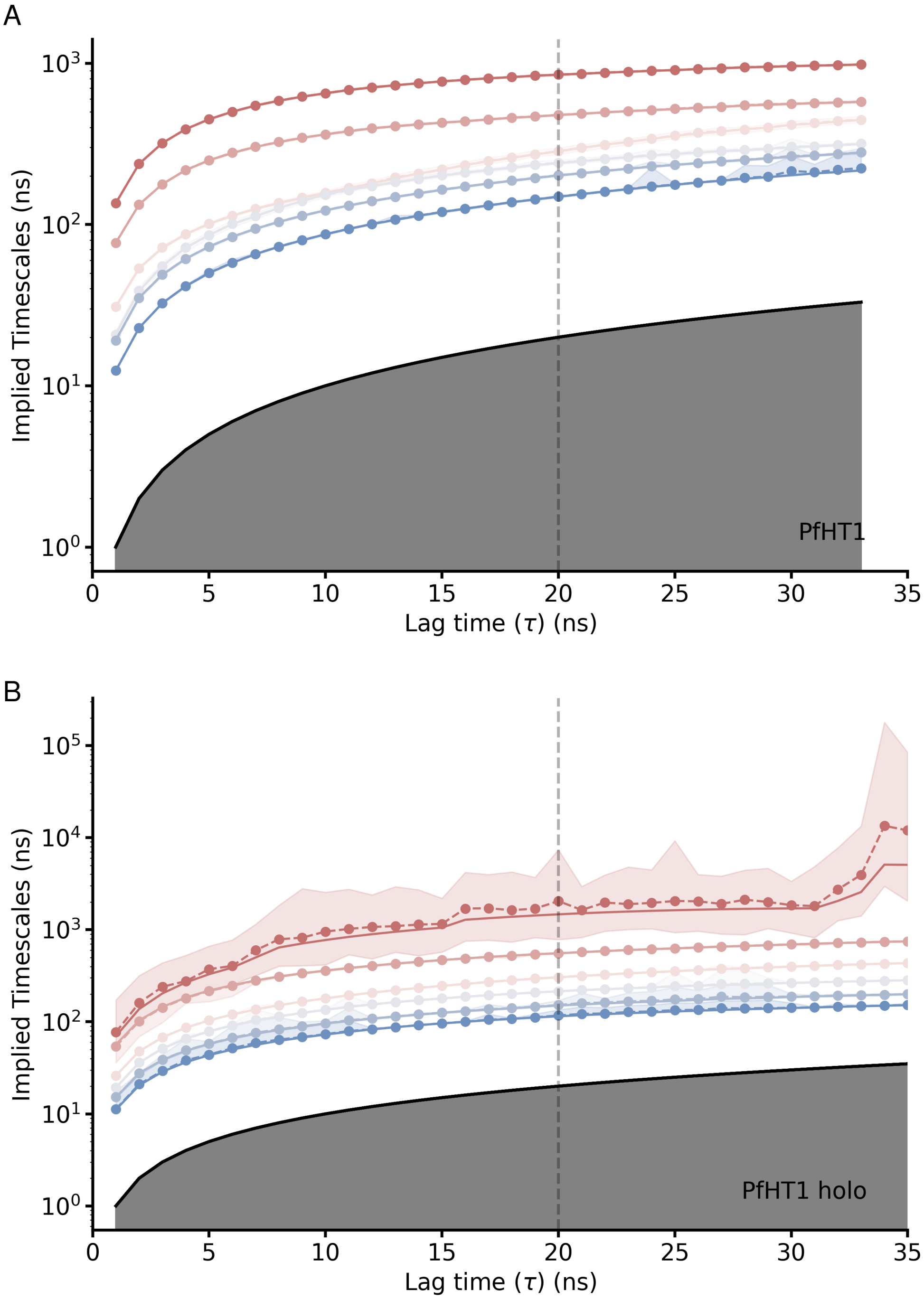


Figure 12: Implied timescale plots for A) Apo systems and B) Holo systems. Based on these plots a lagtime of 20ns was used as the lagtime for MSM construction in both systems.

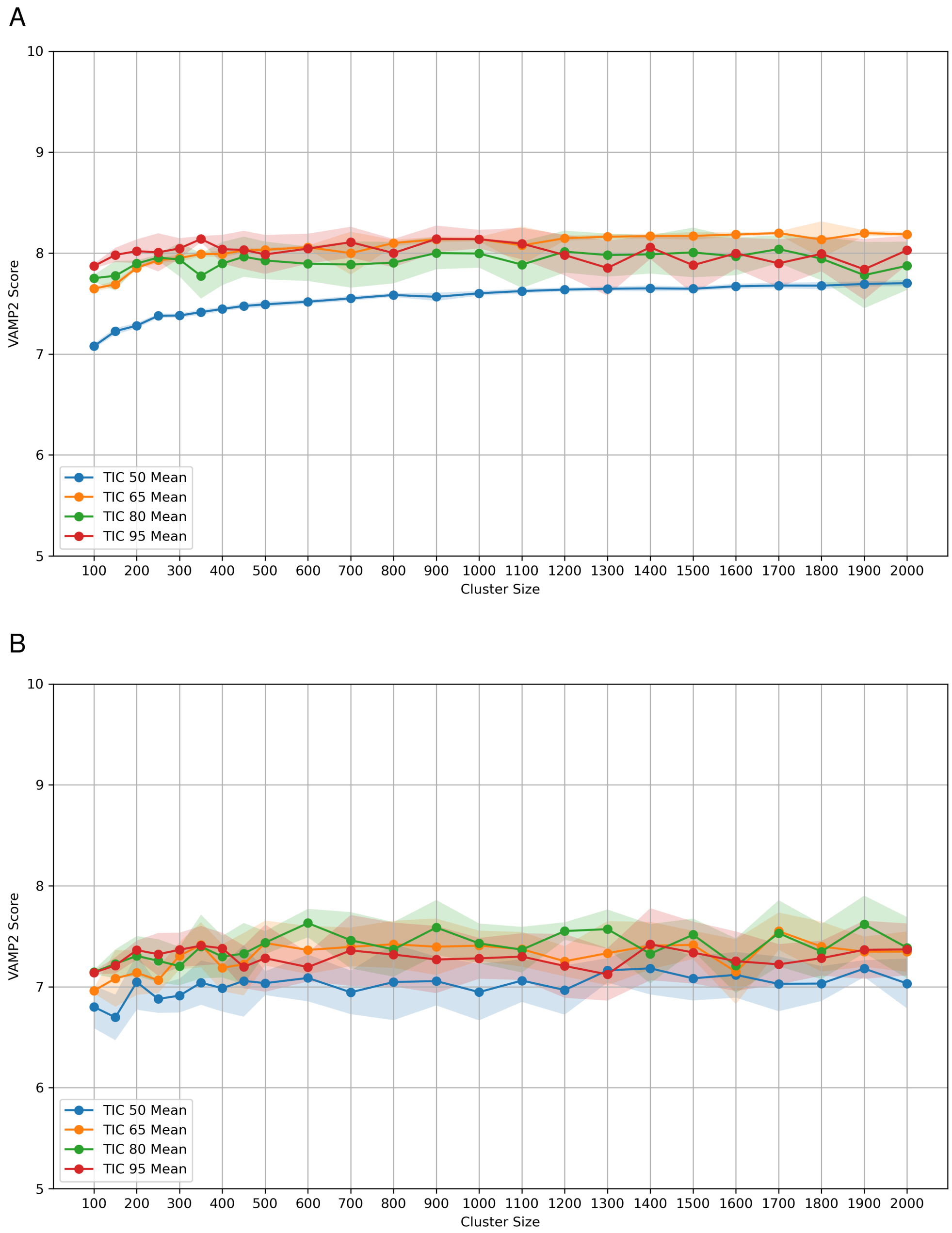


Figure 13: VAMP2 score versus Number of Clusters used for clustering the TICA-reduced data at four different variational cutoffs for A) apo system and B)holo system. The final MSM for apo system was made with 1700 clusters and 65% cutoff and 900 clusters and 80% cutoff.

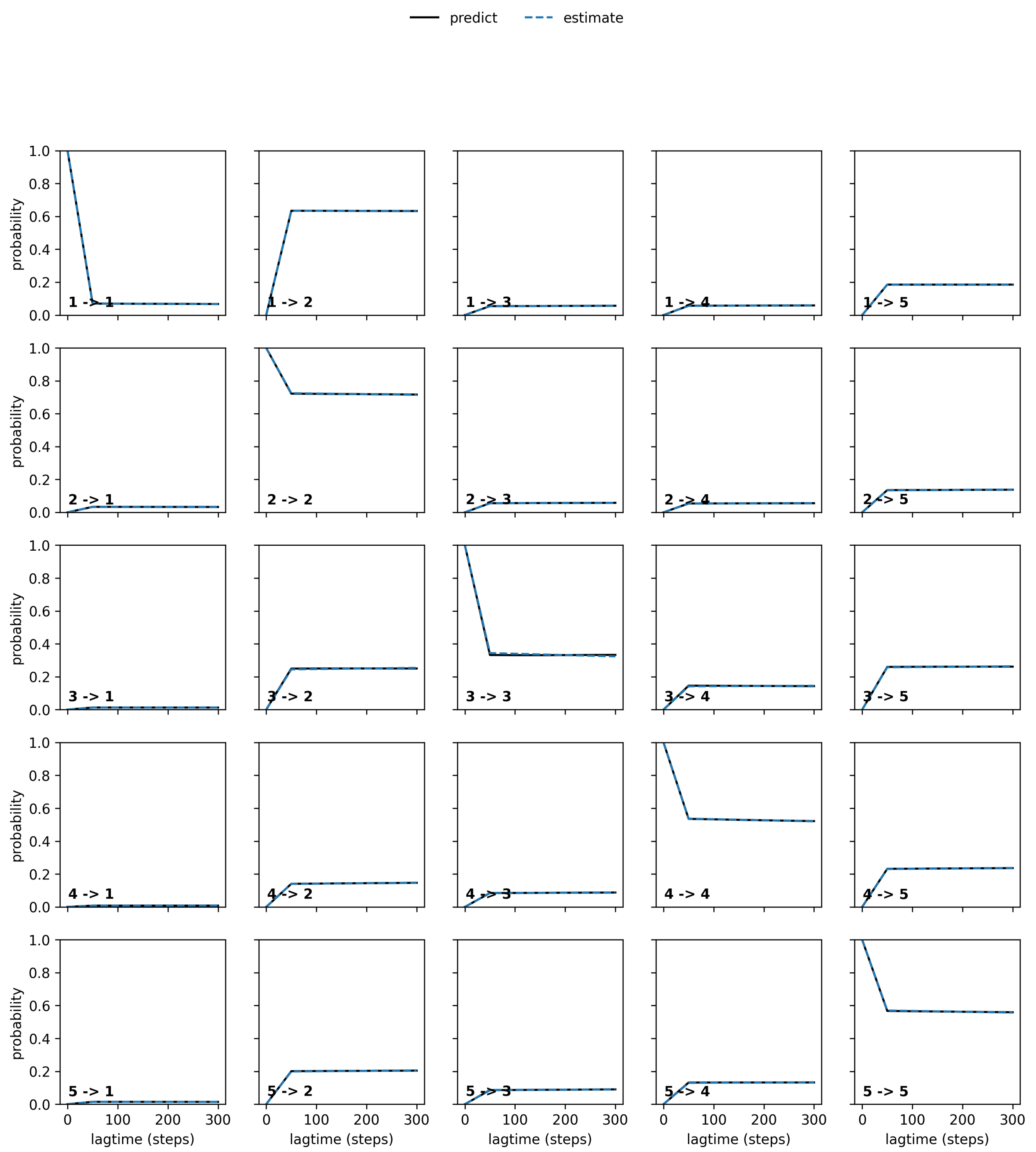


Figure 14: Chapman Kolmogorov test for MSM validation for apo simulations.

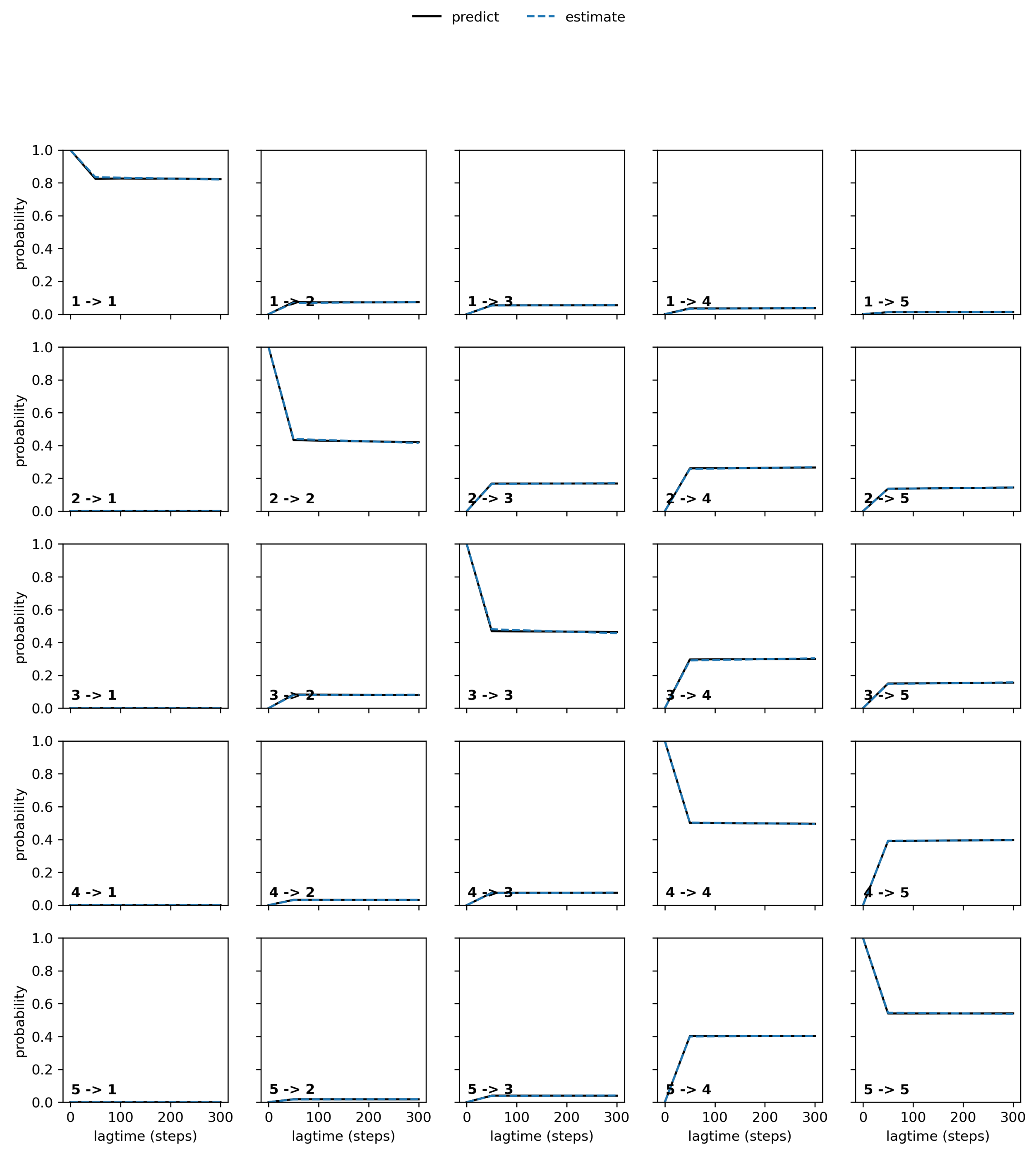


Figure 15: Chapman Kolmogorov test for MSM validation for holo simulations.

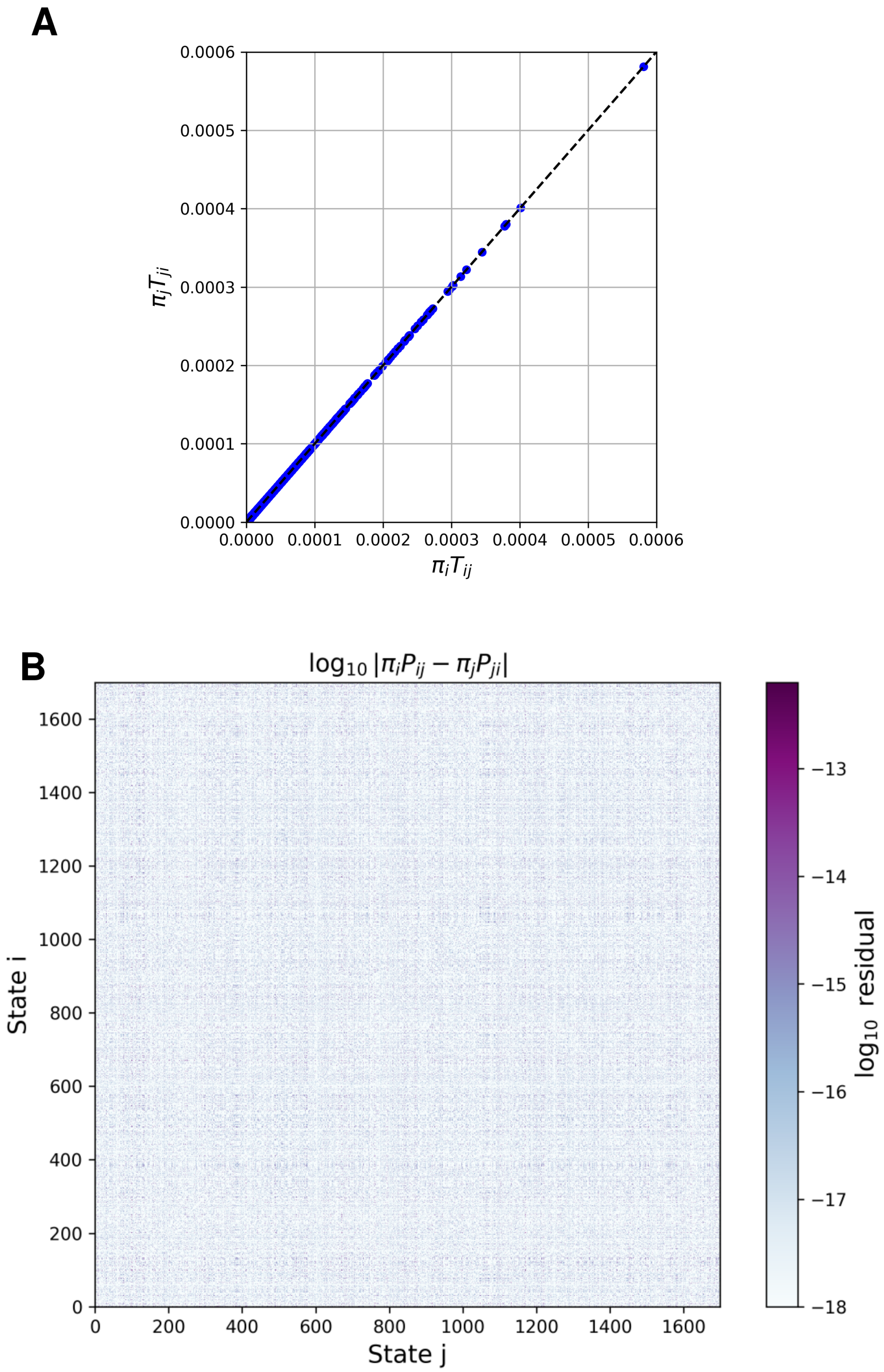


Figure 16: Reversibility of MSM transition probability matrix for apo system. The reversibility was assessed if condition $\pi_i P_{ij} = \pi_j P_{ji}$ held for all MSM states i and j. (A) he quantities $\pi_i P_{ij}$ and $\pi_j P_{ji}$ plotted against each other for all MSM states. (B) The lof of difference between $\pi_i P_{ij}$ and $\pi_j P_{ji}$ showing the two quantities vary within $10^{-13}$.

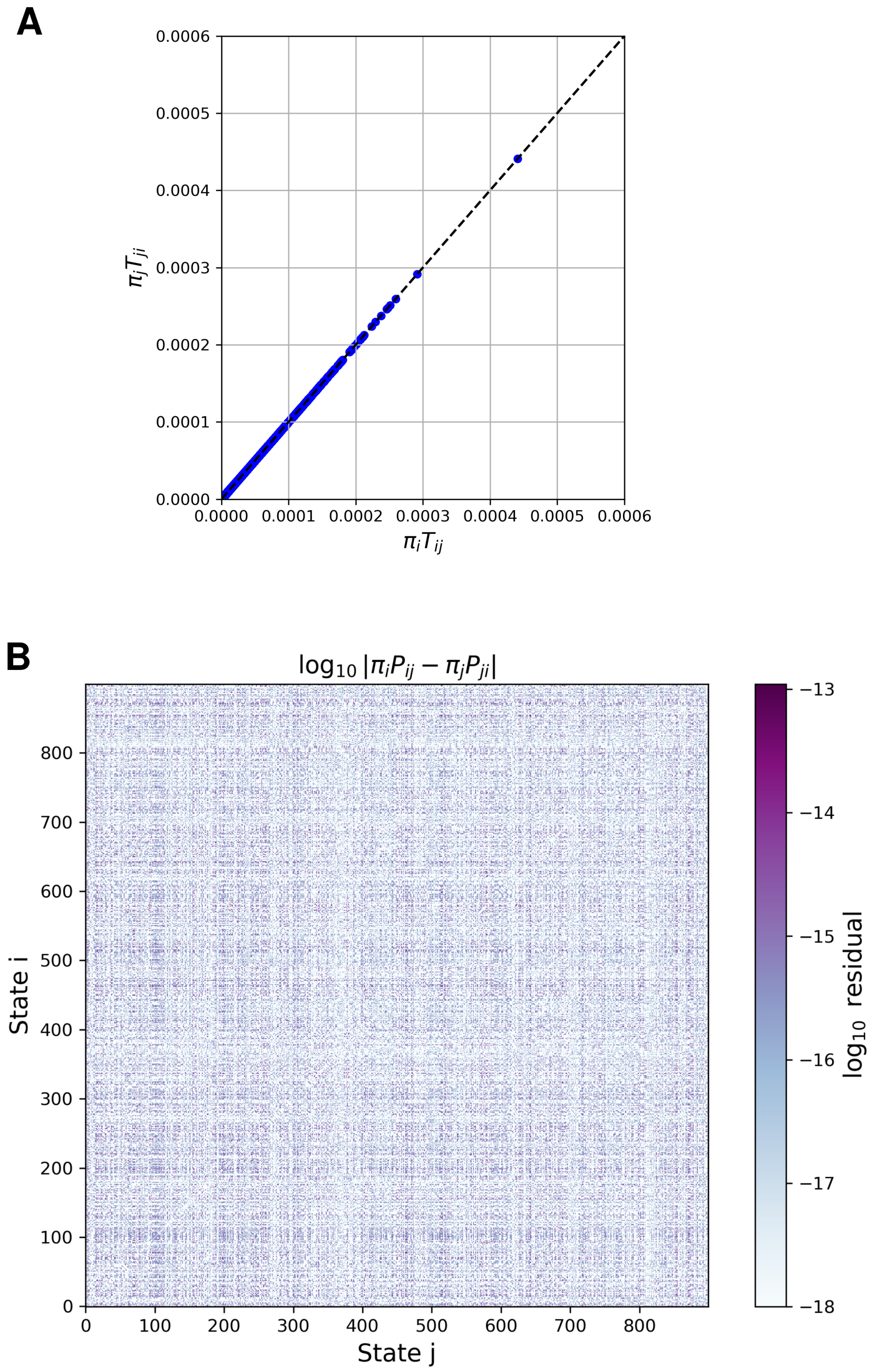


Figure 17: Reversibility of MSM transition probability matrix for holo system. The reversibility was assessed if condition $\pi_i P_{ij} = \pi_j P_{ji}$ held for all MSM states i and j. (A) The quantities $\pi_i P_{ij}$ and $\pi_j P_{ji}$ plotted against each other for all MSM states. (B) The lof of difference between $\pi_i P_{ij}$ and $\pi_j P_{ji}$ showing the two quantities vary within $10^{-13}$.

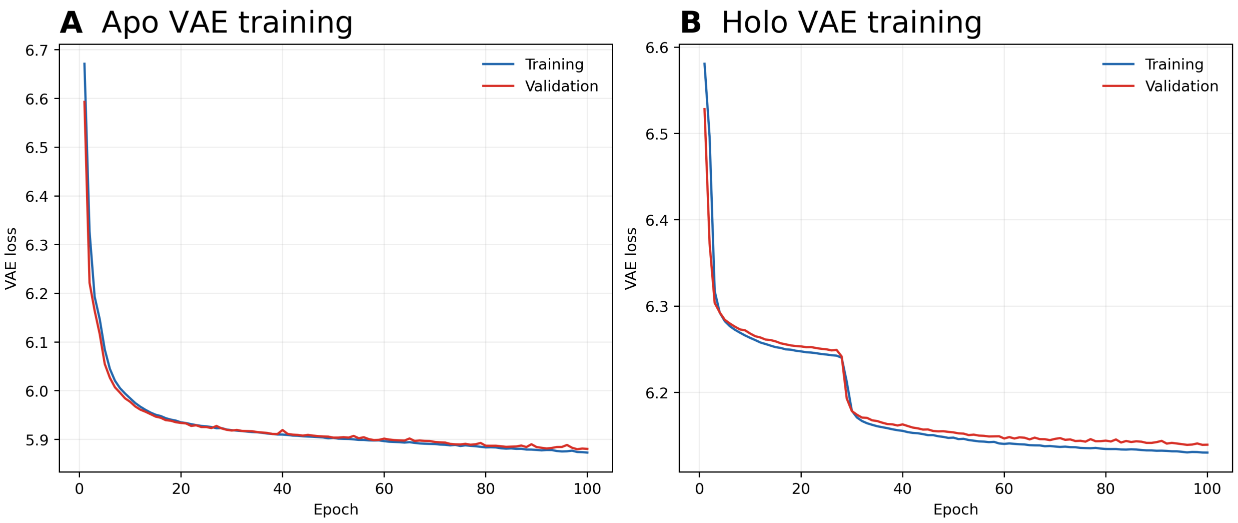


Figure 18: Training and validation losses over 100 epochs for (A) the apo VAE and (B) the glucose-bound VAE. The close training and validation curves show similar optimization behavior for the fitted and held-out pathway subsets.

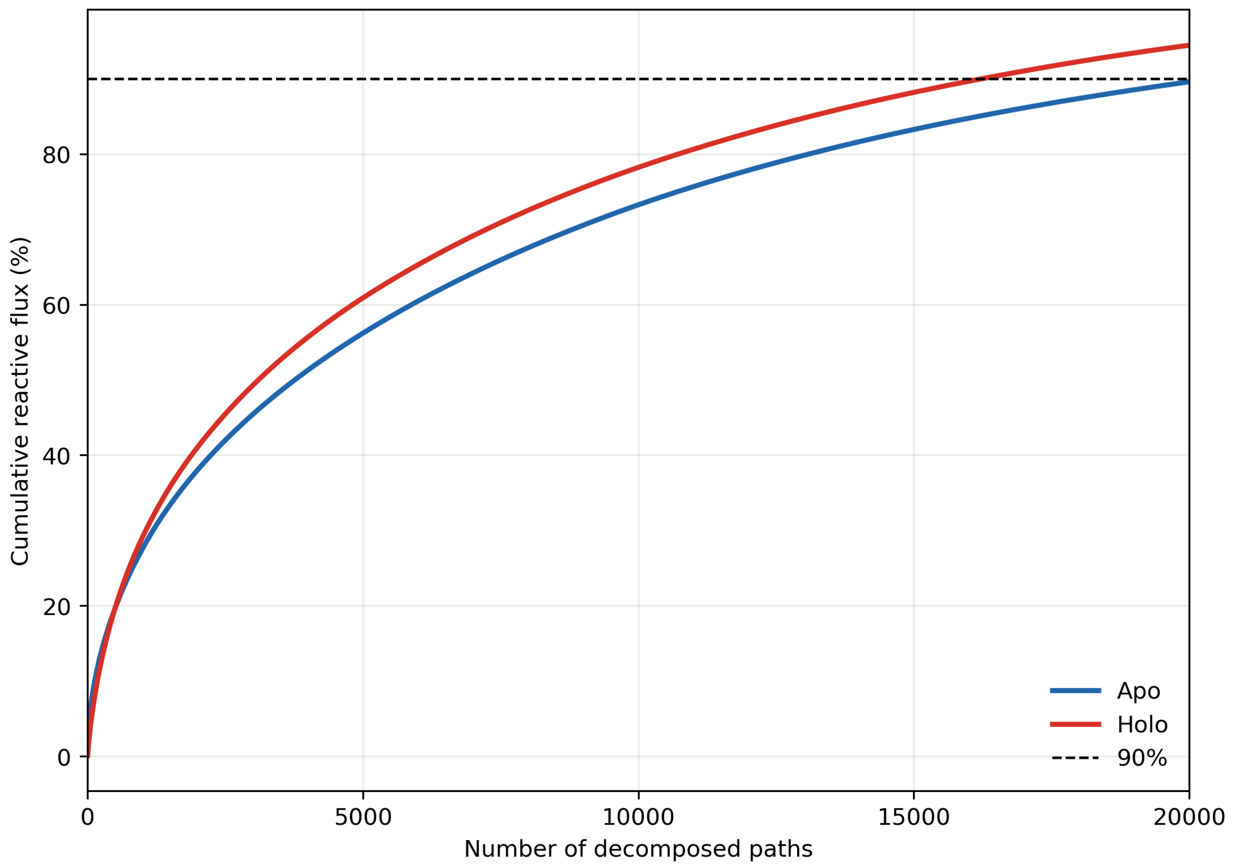


Figure 19: Candidate solutions with one to six K-means clusters are compared using (A) within-cluster sum of squares, (B) silhouette coefficient and (C) gap statistic. Two clusters were selected for both apo and holo, however holo data supports the trivial one cluster solution based on gap statistic.

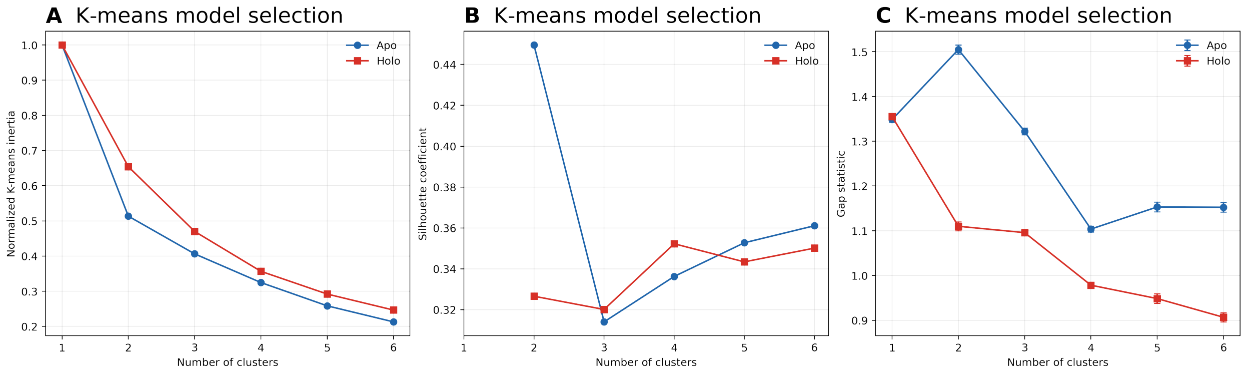


Figure 20: Cumulative fraction of total OF to IF reactive flux recovered as TPT pathways are added in decreasing-flux order for apo and glucose-bound PfHT1. Markers show 5,000, 10,000, and 20,000 paths and the dashed line marks 90% of the total reactive flux.

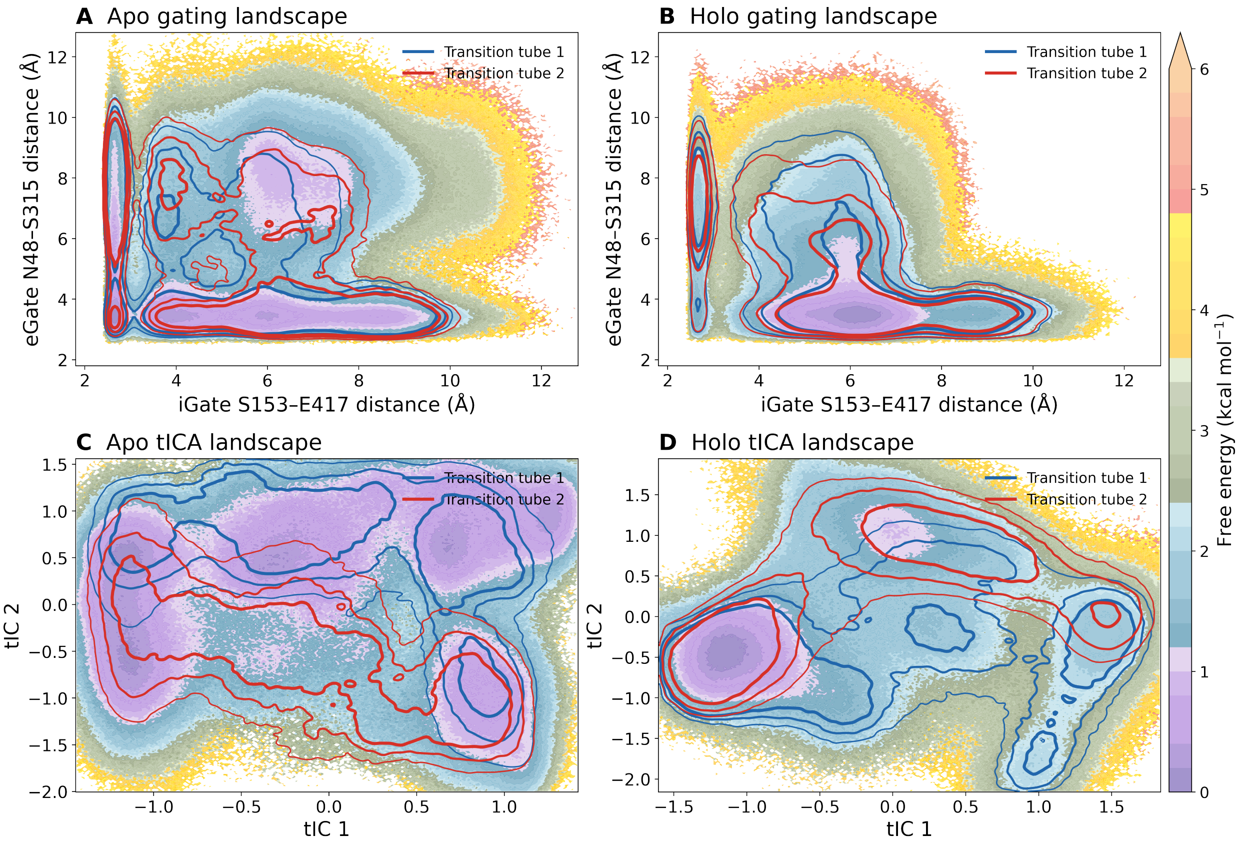


Figure 21: Cluster transition tubes projected onto the MSM-reweighted gating landscapes for (A) apo and (B) holo. The same tubes projected onto the independently estimated tICA landscapes for (C) apo and (D) holo. Contours enclose 50%, 75%, and 90% of each cluster's flux-weighted state density. Background free energies are masked where no trajectory frames were observed.

Table 1: Adaptive sampling statistics for apo and holo PfHT1 simulations.

| **Apo Round** | **Sampling ($\mu$s)** | **# Traj** | **Holo Round** | **Sampling ($\mu$s)** | **# Traj** |
|---|---|---|---|---|---|
| 1 | 25 | 50 | 1 | 20 | 200 |
| 2 | 5 | 50 | 2 | 20 | 200 |
| 3 | 5 | 50 | 3 | 20 | 200 |
| 4 | 5 | 50 | 4 | 20 | 200 |
| 5 | 20 | 200 | 5 | 20 | 200 |
| 6 | 20 | 200 | 6 | 20 | 200 |
| 7 | 20 | 200 | 7 | 20 | 200 |
| 8 | 20 | 200 | 8 | 20 | 200 |
| 9 | 20 | 200 | 9 | 20 | 200 |
| 10 | 20 | 200 | 10 | 20 | 200 |
| 11 | 20 | 200 | 11 | 20 | 200 |
| 12 | 20 | 200 | 12 | 20 | 200 |
| 13 | 20 | 200 | 13 | 20 | 200 |
| 14 | 20 | 200 | 14 | 20 | 200 |
| 15 | 20 | 200 | 15 | 20 | 200 |
| 16 | 20 | 200 | 16 | 20 | 200 |
| 17 | 20 | 200 | 17 | 20 | 200 |
| 18 | 20 | 200 | 18 | 20 | 200 |
| 19 | 20 | 200 | 19 | 20 | 200 |
| 20 | 20 | 200 | 20 | 20 | 200 |
| 21 | 20 | 200 | 21 | 20 | 200 |
| – | – | – | 22 | 20 | 200 |
| – | – | – | 23 | 20 | 200 |
| **Total** | **380$\mu$s** | **3600** | | **460$\mu$s** | **4600** |

Table 2: Membrane composition used for PfHT1 simulations. Lipid species are grouped by class with corresponding counts.

| **Category** | **Lipid Species** | **Count** |
|---|---|---|
| Cholesterol | (CHOL) | 62 |
| 16:0/18:0 | (PSPC) | 34 |
| 16:0/18:1 | (POPC) | 46 |
| 18:2/18:2 | (DLiPC) | 10 |
| 16:0/18:0 | (PSPE) | 22 |
| 16:0/18:1 | (POPE) | 28 |
| 18:2/18:2 | (DLiPE) | 6 |
| 16:0/18:0 | (PSPS) | 10 |
| 16:0/18:1 | (POPS) | 18 |
| 16:0/18:0 | (PSPI) | 4 |
| 16:0/18:1 | (POPI) | 4 |
| 16:0/18:0 | (PSPA) | 6 |
| 16:0/18:1 | (POPA) | 6 |

Table 3: Structural features used for MSM construction. Residue-pair distances were calculated using the closest-heavy atom scheme.

| **Feature Type** | **Residue** | **Residue** |
|---|---|---|
| Distance | Leu182 | Val336 |
| Distance | Ile75 | Lys452 |
| Distance | Leu90 | Ser434 |
| Distance | Gly87 | Ile441 |
| Distance | Val178 | Val340 |
| Distance | Leu80 | Pro445 |
| Distance | Glu76 | Pro445 |
| Distance | Ser83 | Ile441 |
| Distance | Lys358 | Met418 |
| Distance | Val166 | Tyr414 |
| Distance | Ser315 | Ile401 |
| Distance | Ser315 | Phe177 |
| Distance | Tyr151 | Ser429 |
| Distance | Asn48 | Ser315 |
| Distance | Ser153 | Glu417 |
| TM7b kink angle | COM(305-307) - COM(312-314) - COM(320-322) | |

Table 4: Pathway variational autoencoder architecture and training hyperparameters. A single trained model was used for each biochemical condition.

| Parameter | Value |
|---|---|
| Pathway representation | Normalized $30 \times 30$ occupancy histogram in the first two kinetic-map-scaled tICs (900 input values) |
| Encoder architecture | $900 \rightarrow 256 \rightarrow 64$ |
| Latent representation | Two-dimensional Gaussian latent variable represented by encoder means and log-variances |
| Decoder architecture | $2 \rightarrow 64 \rightarrow 256 \rightarrow 900$, followed by a log-softmax output |
| Training objective | Categorical cross-entropy reconstruction loss plus $\beta D_{\mathrm{KL}}$ |
| $\beta$ | 0.01 |
| KL warmup | Linear warmup over 30 epochs |
| Training epochs | 100 |
| Batch size | 256 pathways |
| Optimizer learning rate | $1.0 \times 10^{-3}$ |
| Validation fraction | 0.20 |
| Number of pathways | 20,000 per biochemical condition |
| VAE realization used | Apo: seed 167; glucose-bound: seed 239 |
| Clustering input | Two-dimensional VAE encoder means |
| K-means parameters | $k = 2$, Lloyd algorithm, 100 centroid initializations, random state 20260730 |

Table 5: Structural features associated with tIC2 and transition-tube separation. $r_{\mathrm{tIC2}}$ is the Pearson correlation between each input feature and tIC2, while $\beta_{\mathrm{std}}$ is its standardized coefficient in the linear tICA transformation. Tube averages were calculated using flux-weighted, path-occupancy-normalized microstate populations. Distance values are in Å; TM7b values are in degrees. $\Delta = \text{Tube 2} - \text{Tube 1}$. The sign of tIC2 is arbitrary and should not be compared between the separately fitted apo and glucose-bound models.

| System | Feature | $r_{\mathrm{tIC2}}$ | $\beta_{\mathrm{std}}$ | Tube 1 | Tube 2 | $\Delta$ |
|---|---|---|---|---|---|---|
| Apo | L80–P445 | 0.921 | 0.532 | 13.18 | 11.29 | -1.89 |
| | I75–K452 | 0.706 | 0.046 | 13.23 | 10.86 | -2.37 |
| | Q76–P445 | 0.896 | 0.109 | 13.56 | 12.00 | -1.56 |
| | V166–Y414 | 0.324 | 0.120 | 9.78 | 9.20 | -0.58 |
| | S83–I441 | 0.591 | -0.007 | 11.47 | 11.17 | -0.30 |
| | eGate (N48–S315) | 0.088 | -0.091 | 5.36 | 5.47 | 0.10 |
| | iGate (S153–E417) | -0.753 | -0.338 | 5.64 | 6.11 | 0.47 |
| | TM7b angle | 0.058 | -0.038 | 128.03 | 133.56 | 5.53 |
| Glucose-bound | L80–P445 | 0.744 | 0.400 | 11.54 | 12.53 | 1.00 |
| | I75–K452 | 0.472 | 0.018 | 11.59 | 12.61 | 1.02 |
| | Q76–P445 | 0.719 | 0.420 | 11.61 | 12.50 | 0.89 |
| | G87–I441 | 0.587 | 0.063 | 11.95 | 12.18 | 0.23 |
| | S83–I441 | 0.623 | 0.133 | 10.80 | 11.10 | 0.30 |
| | eGate (N48–S315) | -0.101 | -0.078 | 5.17 | 5.01 | -0.16 |
| | iGate (S153–E417) | -0.228 | -0.087 | 5.97 | 5.86 | -0.11 |
| | TM7b angle | -0.264 | -0.777 | 134.00 | 129.33 | -4.67 |

### Supplementary Methods: Transition-pathway analysis

#### Transition-path theory and pathway decomposition

Transition-path theory (TPT) was applied separately to the equilibrium transition matrix, $\mathbf{T}$, and stationary distribution, $\boldsymbol{\pi}$, of the apo and glucose-bound PfHT1 Markov state models (MSMs). For source ensemble $A$ and target ensemble $B$, the gross reactive current from microstate $i$ to microstate $j$ was

$$f_{ij} = \pi_i q_i^- T_{ij} q_j^+, \tag{1}$$

where $q_i^+$ and $q_i^-$ are the forward and backward committors, respectively. The positive net reactive current was

$$f_{ij}^+ = \max(0, f_{ij} - f_{ji}) \, . \tag{2}$$

Outward-facing (OF) and inward-facing (IF) endpoint ensembles were defined using the mean gate distances of each microstate and the conformational ranges used in Fig. 1 of the main text. The OF-to-IF net current was decomposed using an iterative widest-path procedure. At each iteration, the source-to-target path with the largest bottleneck current was identified. The minimum residual edge current along that path was assigned as its pathway flux and subtracted from every edge in the path before the next iteration. Paths were retained in decreasing-flux order. The first 20,000 paths captured 89.61% of the total apo reactive flux and 94.47% of the total glucose-bound reactive flux. All subsequent pathway embedding, clustering, gate-order, and transition-tube analyses used these 20,000 explicitly decomposed paths.

#### Pathway fingerprints

Each pathway was represented by its occupancy in the first two kinetic-map-scaled tICA coordinates. Because the apo and glucose-bound tICA models were fitted independently, fingerprint construction and all subsequent latent-space analyses were performed separately for the two systems; their tICA and latent axes were not compared numerically. For each system, the tICA range was bounded by the 0.5th and 99.5th percentiles of the sampled coordinates and divided into a 30 × 30 grid. To reduce domination by highly sampled equilibrium regions, each tICA bin $b$ was assigned the inverse-density weight

$$w_b = \min\left[10,\ \frac{\mathrm{median}_{b':C_{b'}>0}\, C_{b'}}{C_b + 1}\right], \tag{3}$$

where $C_b$ is the number of sampled frames in bin $b$. These weights were used to construct a density-corrected histogram, $H_{sb}$, for each microstate $s$. For pathway $p$, with ordered microstate sequence $\mathcal{S}_p$, the fingerprint was

$$x_{pb} = \frac{\displaystyle\sum_{s\in\mathcal{S}_p} H_{sb}}{\displaystyle\sum_{b'}\sum_{s\in\mathcal{S}_p} H_{sb'}}. \tag{4}$$

Every pathway fingerprint therefore had unit $L_1$ mass. Endpoint microstates were retained. Pathway flux was stored separately and was not used as a VAE input or training weight.

#### Variational autoencoder

A separate variational autoencoder (VAE) was trained for each system to embed the 900-dimensional pathway fingerprints into two dimensions. The encoder consisted of fully connected layers containing 256 and 64 nodes, followed by two-dimensional latent mean and log-variance layers. The decoder contained layers of 64 and 256 nodes followed by a log-softmax output over the 900 fingerprint bins. For input fingerprint $\mathbf{x}_p$, encoder $q_\varphi(\mathbf{z}_p \mid \mathbf{x}_p)$, and decoder $p_\theta(\mathbf{x}_p \mid \mathbf{z}_p)$, the loss was

$$\mathcal{L} = -\sum_p\sum_b x_{pb}\log\widehat{x}_{pb} + \beta\sum_p D_{\mathrm{KL}}[q_\varphi(\mathbf{z}_p \mid \mathbf{x}_p)\,\|\,\mathcal{N}(\mathbf{0},\mathbf{I})], \tag{5}$$

where $\widehat{x}_{pb}$ is the reconstructed bin probability and $\beta = 0.01$. The KL contribution was introduced linearly over the first 30 epochs.

Each model was trained for 100 epochs using the Adam optimizer, a batch size of 256, and a learning rate of $10^{-3}$. Pathways were divided into 80% training and 20% validation subsets using a group-aware random split (random state 20260804), with the SHA-256 hash of the ordered microstate sequence used as the group identifier. The checkpoint with the lowest validation loss was retained. One fixed VAE realization was used for each system: seed 167 for apo and seed 239 for glucose-bound PfHT1. Training and validation loss curves were inspected to assess optimization and overfitting. The posterior means of all 20,000 pathways were used as the two-dimensional clustering coordinates.

#### K-means clustering and cluster-number evaluation

The apo and glucose-bound latent representations were clustered separately using K-means with the Lloyd algorithm. The final analysis used two clusters, 100 centroid initializations, and random state 20260804. TPT flux was not used during clustering. After fitting, cluster numbers were assigned in decreasing order of the summed TPT flux of their constituent pathways and were reported as Cluster 1 and Cluster 2. Two clusters were chosen for apo and holo latent spaces using the within-cluster sum of squares, silhouette coefficient, and gap statistic. The corresponding projected pathway ensembles are referred to as Transition tube 1 and Transition tube 2. For apo PfHT1, the two-cluster silhouette coefficient was 0.450, compared with 0.314 for three clusters, and the unrestricted gap statistic selected two clusters. For glucose-bound PfHT1, the two-cluster silhouette coefficient was 0.327, compared with 0.320 for three clusters, and the gap statistic selected two when comparison was restricted to nontrivial partitions containing at least two clusters. However, the unrestricted glucose-bound gap statistic, including the one-cluster model, favored a single diffuse population. The two glucose-bound clusters were therefore retained as a geometric partition of a continuous pathway ensemble and were not interpreted as evidence for two discrete molecular mechanisms.

#### Gate-event classification

Gate-event order was assigned directly from the ordered microstate sequence of each decomposed pathway and independently of its K-means label. For each microstate, gate coordinates were the mean N48–S315 eGate distance and mean S153–E417 iGate distance. The first microstate at which the eGate distance was $\leq 6.25$ Å and the first microstate at which the iGate distance was $\geq 6.75$ Å were identified. A pathway was classified as eGate-closing-first when the eGate event preceded the iGate event, iGate-opening-first when the order was reversed, and concerted when both events first occurred in the same microstate. Gate-order fractions within each latent cluster were calculated by summing the TPT fluxes of its constituent pathways in each category and normalizing by the total flux of that cluster. Flux outside the 20,000 explicitly decomposed paths was not assigned a gate-event category.

#### Transition-tube visualization

Cluster-specific transition tubes were constructed from the explicit pathway ensemble. Each pathway distributed its TPT flux equally among the microstates in its sequence, preventing longer paths from receiving greater weight solely because they contained more states. These contributions were summed separately for Cluster 1 and Cluster 2 and normalized to unit mass. The mass assigned to each microstate was then distributed uniformly over the trajectory frames assigned to that microstate and projected onto either the gating coordinates or the first two tICA coordinates. Projected cluster densities were evaluated on a $250 \times 250$ grid and smoothed using a Gaussian kernel with a width of two bins. Contours were defined as highest-density regions enclosing 50%, 75%, and 90% of the normalized cluster mass. The cluster-specific projected ensembles are referred to as Transition tube 1 and Transition tube 2 and were displayed in blue and red, respectively. Background free-energy surfaces were calculated from equilibrium trajectory weights obtained from the corresponding MSM.

#### Committor projection within transition tubes

The forward committor obtained from the endpoint-specific TPT calculation was used to visualize progress along each transition tube. For microstate $i$, the forward committor

$$q_i^+ = \Pr_i(\tau_{\mathrm{IF}} < \tau_{\mathrm{OF}}) \tag{6}$$

is the probability that a trajectory initiated in microstate $i$ reaches the IF ensemble before returning to the OF ensemble. Consequently, $q_i^+ = 0$ for OF source states and $q_i^+ = 1$ for IF target states. The committor was calculated once from the complete OF-to-IF TPT problem and was not recalculated separately for each latent cluster.

For each transition tube, the through-state weight of microstate $i$ was calculated by summing the fluxes of the cluster-assigned decomposed paths that visited that microstate. Each microstate was placed at its MSM-weighted centroid in the first two tICA coordinates. The state weights and committor-weighted state weights were histogrammed on a $180 \times 180$ grid and smoothed separately using a Gaussian kernel with a width of three bins. The displayed committor field was their ratio,

$$\overline{q}_c^+(x, y) = \frac{G_\sigma\left[\sum_i J_{ci} q_i^+ \delta_{b(i)}(x, y)\right]}{G_\sigma\left[\sum_i J_{ci} \delta_{b(i)}(x, y)\right]}, \tag{7}$$

where $J_{ci}$ is the through-state flux of microstate $i$ in transition tube $c$, $b(i)$ is the tICA bin containing its centroid, and $G_\sigma$ denotes Gaussian smoothing.

The field was restricted to the highest-density support containing 95% of the smoothed tube weight. Contour lines were drawn at forward committor intervals of 0.1. Microstates with nonzero tube-specific through-state flux that fell inside this support were displayed as uniform dark points. The committor field was shown using a semi-transparent two-color scale over the MSM-weighted tICA free-energy landscape. Because interpolation and smoothing were used only for visualization, separately plotted exact microstate committors were retained as a validation of the projected fields.

#### Software and reproducibility

MSM and TPT calculations were performed using `deeptime` 0.4.4. VAE training and the final K-means analysis used PyTorch and `scikit-learn`. The retained analysis package contains the pathway fluxes and microstate sequences, selected latent embeddings, cluster assignments, MSM-weighted landscape coordinates, and all parameters required to reproduce the clustering and visualization.
